# Inflammation promotes tumor aggression by stimulating stromal cell-dependent collagen crosslinking and stromal stiffening

**DOI:** 10.1101/2020.02.13.948141

**Authors:** Ori Maller, Allison P. Drain, Alexander S. Barrett, Signe Borgquist, Brian Ruffell, Pham T Thanh, Tina Gruosso, Hellen Kuasne, Johnathon N. Lakins, Irene Acerbi, J. Matthew Barnes, Travis Nemkov, Aastha Chauhan, Jessica Gruenberg, Aqsa Nasir, Olof Bjarnadottir, Zena Werb, Peter Kabos, E. Shelley Hwang, Morag Park, Lisa M. Coussens, Andrew C. Nelson, Kirk C. Hansen, Valerie M. Weaver

**Affiliations:** Department of Surgery, Center for Bioengineering and Tissue Regeneration, University of California, San Francisco, California, USA; Department of Biochemistry and Molecular Genetics, University of Colorado Denver - Anschutz Medical Campus, Aurora, CO, USA; Division of Oncology and Pathology, Department of Clinical Sciences, Lund, Lund University; Clinical Trial Unit, Clinical Studies Sweden, Forum South, Skåne University Hospital, Lund, Sweden; Cell, Developmental & Cancer Biology, Oregon Health & Science University; Knight Cancer Institute, Oregon Health & Science University, Portland, Oregon, USA; Goodman Cancer Research Centre, McGill University, Montreal, QC, Canada; Department of Biochemistry, McGill University, Montreal, QC, Canada; Department of Oncology, McGill University, Montreal, QC, Canada; Department of Laboratory Medicine and Pathology, University of Minnesota, Minneapolis, MN, USA; Department of Anatomy and Biomedical Sciences Program, University of California, San Francisco CA, USA; UCSF Helen Diller Comprehensive Cancer Center, University of California, San Francisco, San Francisco, CA, USA; Department of Medicine, Division of Medical Oncology, University of Colorado Anschutz Medical Campus, Aurora, Colorado, USA; Department of Pathology, University of California, San Francisco, CA, USA; Department of Surgery, Duke University Medical Center, Durham, NC, USA; Departments of Bioengineering and Therapeutic Sciences, and Radiation Oncology, Eli and Edythe Broad Center of Regeneration Medicine and Stem Cell Research

## Abstract

Collagen deposition and stromal stiffening accompany malignancy, compromise treatment, and promote tumor aggression. Clarifying the molecular nature of and the factors that regulate extracellular matrix stiffening in tumors should identify biomarkers to stratify patients for therapy and therapeutic interventions to improve outcome. We profiled lysyl hydroxylase- and lysyl oxidase-mediated collagen crosslinks and quantified the greatest abundance of total and complex collagen crosslinks in more aggressive human breast cancer subtypes with the stiffest stroma. These tissues also harbored the highest number of tumor-associated macrophages (TAM), whose therapeutic ablation not only reduced metastasis, but also concomitantly decreased accumulation of collagen crosslinks and stromal stiffening. Epithelial-targeted expression of the crosslinking enzyme lysyl oxidase had no impact on collagen crosslinking in PyMT mammary tumors, whereas stromal cell targeting did. Consistently, stromal cells in microdissected human tumors expressed the highest level of collagen crosslinking enzymes. Immunohistochemical analysis of a cohort of breast cancer patient biopsies revealed that stromal expression of lysyl hydroxylase two, an enzyme that induces hydroxylysine aldehyde-derived collagen crosslinks and stromal stiffening correlated significantly disease specific mortality. The findings link tissue inflammation, stromal cell-mediated collagen crosslinking and stiffening to tumor aggression and identify lysyl hydroxylase two as a novel stromal biomarker.

**Significance:** We show infiltrating macrophages induce stromal fibroblast, and not epithelial, expression of collagen crosslinking enzymes that drive tumor stiffening. Stromal enzyme LH2 is significantly upregulated in breast cancer patients with the stiffest stroma, the most trivalent HLCCs and the worst prognosis, underscoring its potential as a biomarker and therapeutic target.

## Introduction

Pathological accumulation of extracellular matrix (ECM) accompanies the formation of all solid tumors(1–3). The tumor ECM is composed primarily of interstitial collagen that is progressively reorganized and stiffened(2, 4). The collagenous fibrotic tumor ECM compromises treatment and is linked to poor patient prognosis(5–8). Tumor biopsy analysis showed that a thick fibrous collagenous ECM associates with less differentiated tumors and that this phenotype predicts poor patient survival, emphasizing the relevance of collagen architecture(1,4,9). Patients with pancreatic ductal adenocarcinomas (PDACs) that are surrounded by stiff, thick fibrous collagens have a shorter survival, and invasive breast carcinomas with the stiffest ECM stroma at their invasive front are the most aggressive(1, 2). These observations suggest that stromal stiffness reflects collagen organization may be an important prognostic variable. Consistently, preclinical studies using organotypic cultures and rodent models provide plausible evidence for a causal relationship between collagen organization, stromal stiffness, tumor cell invasion in culture, and metastasis *in vivo*(10–14). These findings underscore the clinical relevance of collagen architecture and stiffness to malignancy, and emphasize the need to clarify the molecular nature of the collagenous ECM so that new biomarkers can be identified and anti-cancer therapeutics may be developed(1,2,15,16).

Interstitial type I fibrillar collagen is the ECM component that contributes most significantly to the tensile strength of tissue(17). The tensile strength of interstitial collagen depends upon the activities of two major families of enzymes: the lysyl hydroxylases (LH; gene name procollagen-lysine, 2-oxoglutarate 5-dioxygenase or PLOD) and the lysyl oxidases (LOX), which regulate fibrillogenesis of newly synthesized collagen molecules through intermolecular covalent crosslinking(18–21). Fibrotic human tumors express high levels of LOX and LH enzymes(22). Tumor grade and overall patient survival associate with total tissue LOX and PLOD2 mRNA(21,23–25). Pharmacological or antibody-mediated inhibition of LOX in MMTV-Her2/Neu mice or genetic reduction of PLOD2 in subcutaneously-injected lung tumor epithelial cells reduce tissue fibrosis, stromal stiffening and collagen crosslinking and concomitantly decrease tumor incidence and aggression(12, 23). Moreover, elevating LOX or LH2-mediated collagen crosslinking enhances fibrosis and stromal stiffness and promotes malignant transformation and tumor aggression in lung and mammary xenografted tumors(12, 23). These observations suggest that the direct targeting of specific collagen crosslinking enzymes has clinical merit for the treatment of cancer. However, given caveats with recent clinical trials targeting ECM modifiers including suboptimal activity of inhibitory treatments and the risk of off-target effects, strategies designed to interfere with the induction and activation of these crosslinking enzymes offer an attractive alternative(26). Towards this goal, the identification and causal implication of additional factors that regulate the levels and/or activity of collagen crosslinking enzymes has the potential to identify new predictive biomarkers and alternative anti-tumor treatment targets.

Pre-neoplastic lesions are inflamed, and pathological fibrosis correlates with inflammation(27, 28). Chronic inflammation and experimental manipulations that promote inflammation in rodent models induce fibrosis by secreting factors such as metalloproteinases and TGFβ(14,28–31). Furthermore, fibrotic tumors are frequently inflamed, and this inflammation promotes tumor aggression, whereas either inhibiting inflammation or decreasing macrophage infiltration reduce tumor metastasis and enhance anti-tumor treatment(2,27,32–35). Nevertheless, it remains unclear if inflammation promotes tumor progression and aggression by inducing stromal stiffening, and if so, whether this is regulated via epithelial and/or stromal fibroblast-mediated collagen remodeling and crosslinking.

## Results

### xAAA profiling identifies increased levels of collagen crosslinks and stromal stiffness as indicators of breast tumor aggression

To clarify the role of collagen crosslinking in tumor fibrosis we developed a crosslinked amino acid analysis (xAAA) method that enabled the characterization and quantification of specific collagen crosslinks in tissues across a wide range of collagen levels. We utilized solid phase extraction (SPE) enrichment followed by high pH amide hydrophilic chromatography (HILIC) coupled to a benchtop orbitrap (QExactive) mass spectrometer for these measurements. The validated method detected all known LOX-generated crosslinks including divalent (lysinonorleucine, dihydroxy lysinonorleucine), trivalent (pyridinoline and deoxy-pyridinoline) and tetravalent (desmosine and isodesmosine) crosslinked amino acids linearly over four orders of magnitude, with calculated limits of quantification (LLOQ) in the femtomolar range (**Suppl. Fig. 1; Suppl. Table 1**)(36). The technique revealed a positive correlation between collagen crosslinking and abundance in excised human clinical specimens with very low to very high collagen concentrations and varying mechanical properties (**Suppl. Fig 2**). The method also identified a subset of hydroxylysine aldehyde (Hyl^ald^)-derived collagen crosslinks (HLCCs) crucial for the mechanical strength of tissue(17, 37).

We next obtained snap-frozen biospecimens of normal human breast tissue (N=10; age between 22 and 58) and human tumor biopsies representing early stage (stage 1-2) invasive breast cancers (IBC) excised from mastectomy specimens. Molecular subtyping that subdivides human breast tumors stratified by estrogen receptor (ER+) and human epidermal growth factor receptor two (HER2+) status and ER/PR/HER2-negative (triple negative; TN) is a key determinant used to direct the treatment of breast cancer patients. Accordingly, we chose human breast tumor biopsies that represented ER+ (N=8; age between 42 and 71); HER2+ (N=6; age between 40 and 76) and TN (N=6; age between 50 and 71). H&E stained tissue sections confirmed the presence of normal glandular structures in the normal controls and invasive breast cancer in the tumor specimens (**Fig. 1a; top panels**). Polarized light imaging of Picrosirius-stained (PS) tissue revealed that the normal breast tissue stroma had very little fibrillar collagen, whereas stromal tissue in all patients with IBCs contained abundant fibrillar collagen (**Fig. 1a; middle panels)** that second harmonic generation (SHG) imaging indicated was thicker and more linearized (**Fig. 1a; bottom panels**). Polarized light microscopy and two-photon imaging further revealed that the level of fibrosis in the tissue was higher in HER2+ as compared to the ER+ breast tumors, and was further increased in the TN tumors, consistent with our previous report that TN tumors contain a high density of aligned collagen fibers (**Fig. 1a; middle** and **bottom panels**)(2). AFM microindentation revealed a significant increase in the elastic modulus of the stroma associated with the invasive front of all the IBC tissues (**Fig.1b**)(2,4,12). xAAA analysis revealed a significant increase in total collagen crosslinking in all the IBCs (**Fig. 1c, Suppl. Fig. 5**). These findings are consistent with an association between collagen crosslinks, tissue fibrosis and stromal stiffness, as has been previously documented in experimental murine models of mammary cancer(12). Interestingly, when we subdivided the IBC collagen crosslinking analysis into breast tumor subtype the most significant increase in total collagen crosslinks was calculated to be in the TN breast tumors (**Fig. 1i**). Furthermore, biochemical quantification of total tissue collagen 1A1 or 1A2 did not account for the higher total number of collagen crosslinks in the TN breast tumor tissue (**Fig. 1k**). Instead, molecular characterization of the isolated tissue collagen revealed that the TN tumors had a distinctive crosslink profile due to a strong preference for a combination of DHLNL, Pyr, and d-Pyr crosslinks **(Fig. 1d-j**). Subtype analysis further revealed that the level of hydroxylysine aldehyde-derived collagen crosslinks (HLCCs) in the TN subtype correlated significantly and positively with the stiffness of the stroma at the invasive front of the tumor tissue (**Fig. 1l**). These findings highlight the importance of HLCC collagen crosslinking in breast cancer aggression.

**Figure 1:**
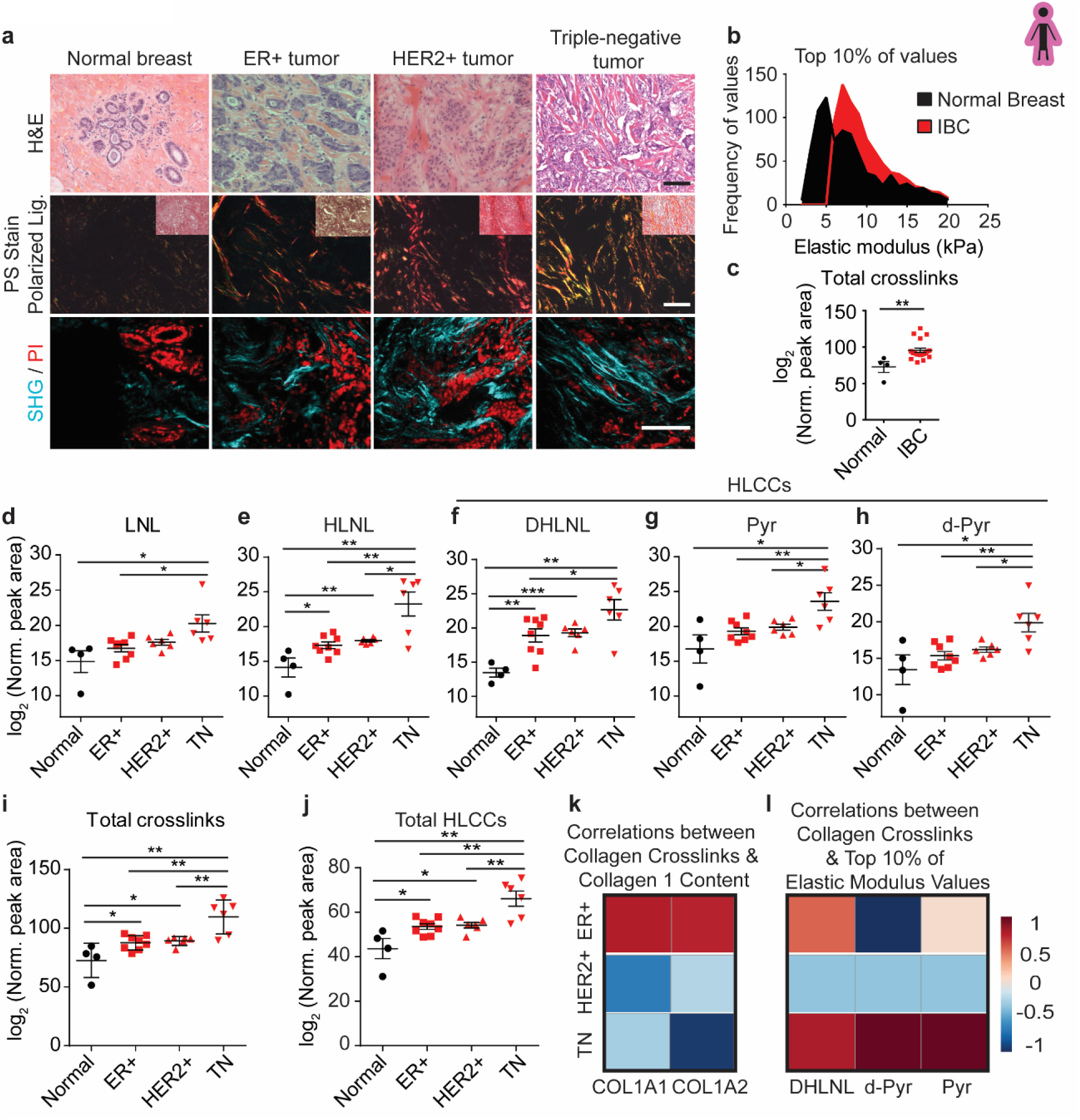
Hydroxylysine collagen crosslink abundance correlates with human breast cancer aggression. (**a**) Representative images of normal breast tissue (normal breast; *n* = 4) and invasive tumors diagnosed as estrogen receptor positive (ER+ tumor; *n* = 8), epidermal growth factor receptor two positive (HER2+ tumor; *n* = 6) and triple negative (TN tumor; *n* = 6). (**Top row**) Brightfield images of human breast tissue stained with hematoxylin and eosin (H&E). (**Middle row**) Polarized light images of picrosirius red (PS) stained human breast tissue with an inset brightfield image shows relative levels of fibrillar collagen. (**Bottom row**) Two photon second harmonic generation (SHG) images of human breast tissue revealing collagen organization (turquoise) and propidium iodide (PI; red) stained nuclei. Scale bar, 100 µm. (**b**) The distribution of the top 10% of elastic modulus values of normal breast tissue (n = 10) and invasive breast carcinoma (IBC; n = 10) measured by AFM microindentation. Statistical analysis was performing using Mann-Whitney U test (****p < 0.0001). (**c**) Quantification of the abundance of all detected collagen crosslinks in normal breast tissue (n = 4) and in IBC tissues (n = 19) plotted as a scatter plot of individual samples with mean ± SEM. Statistical analysis was performed using Mann-Whitney U test (**p < 0.01). (**d-h**) Scatter plots showing individual and mean values ± SEM of the levels of each LCC and HLCC crosslink measured in normal breast tissues, and in ER+, HER2+ and TN breast tumors. The total abundance of crosslinks (**i**) was calculated by summing all individual crosslinks and the total tissue HLCC abundance (**j**) was calculated by summing DHLNL, Pyr, and d-Pyr and plotted as individual and mean values ± SEM. All crosslink values are normalized to total collagen content (i.e. hydroxyproline abundance) and wet tissue weight and are plotted as log_2_ transformed normalized peak areas from LC-MS data. Statistical analysis of crosslinks was performed using one-way ANOVA test for overall analysis and unpaired t-test was used for individual comparisons (*\*p* < 0.05; *\*\*p* < 0.01). (**k,l**) Heat maps of Spearman correlation coefficients indicating correlations between levels of total collagen crosslinks and collagen I content (**k**) and between levels of each HLCC and the top 10% of elastic modulus measurements (**l**) stratified by tumor subtype.

### Increased collagen crosslinking correlates with high expression of stromal LOX and PLOD2 in aggressive tumor subtypes

To determine if the observed increase in collagen crosslinking and stromal stiffness in TN tumors was related to the expression levels of enzymes implicated in regulating collagen crosslinking, we analyzed publicly available human breast cancer gene expression array data (n= 1904) for the genes coding these enzymes and examined their correlation to breast cancer subtype. Bioinformatics analyses revealed a significant increase in the major collagen crosslinking enzyme LOX, but not lysyl oxidase like two (LOXL2), in the more aggressive HER2 and TN tumor subtypes and indicated that LOX levels were particularly high in TN tumors (**Fig. 2a-b**). The arrays also showed bulk gene expression of PLOD2, the major regulator of HLCC accumulation, to be highly upregulated in aggressive human TN breast cancers (**Fig. 2c**). To gain insight into the cellular sources of LOX and LH2 in human tumors, we used laser capture microdissection to isolate regions of tumor epithelium and stroma to identify the origins of LOX and LH2 in invasive human breast cancers (38, 39). Gene expression analysis of stromal and epithelial compartments revealed that the stromal cells in the tumor tissue expressed significantly more LOX and PLOD2 than the associated tumor epithelium. The data further indicated that this relationship was more evident in the breast tissue from women with ER-/PR-breast cancer that is frequently the more aggressive tumor subclass (**Fig. 2d-g**).

**Figure 2:**
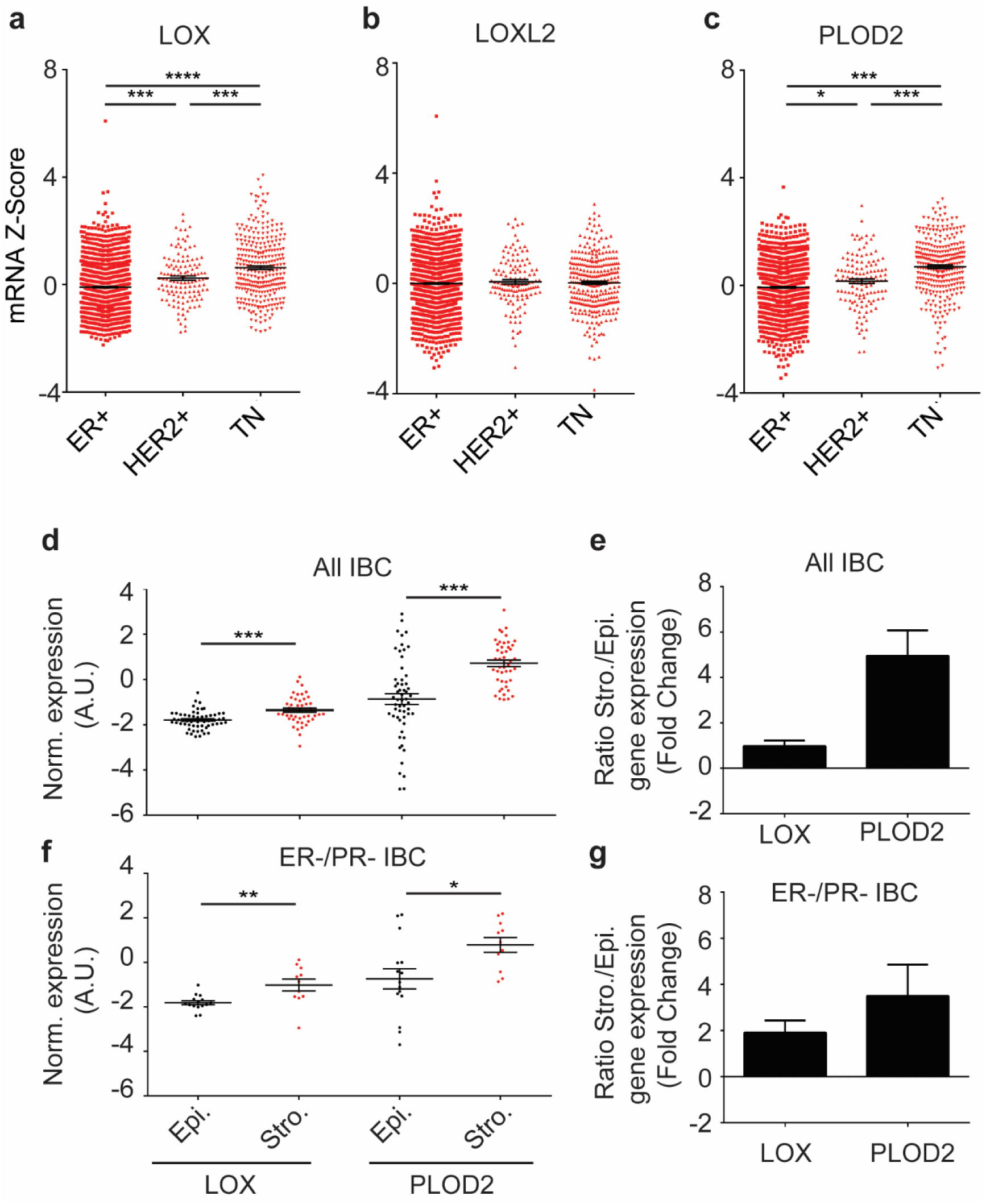
LOX and PLOD2 are enriched in TNBC and predominantly expressed by stromal cells. (**a-c**) Gene expression analysis of LOX (**a**), LOXL2 (**b**), and PLOD2 (**c**) stratified by ER^+^ (n = 1355), HER2^+^ (n = 127), and triple negative (TN; n = 299) subtypes. Gene expression is plotted as a scatter plot of mRNA z scores with the mean ± SEM. Statistical analysis was performed using one-way ANOVA for overall analysis and unpaired t-test was used for individual comparisons (*\*p* < 0.05, **p < 0.01, ***p < 0.001, ****p<0.0001). (**d**) Scatter plot of individual and mean values ± SEM comparing LOX (n = 47) and PLOD2 (n = 57) gene expression in microdissected epithelial and stromal compartments of human invasive breast carcinomas. Statistical analysis was performed using Mann-Whitney U test (****p < 0.0001). (**e**) Quantification of LOX and PLOD2 gene expression fold change from (**d**) in stromal cells relative to epithelial cells. (**f-g**) Restriction of the stromal/epithelial gene expression analysis in (**d**) and (**e**) to estrogen receptor (ER) negative and progesterone receptor (PR) negative samples. (LOX n = 11, PLOD2 n = 15). Statistical analysis was performed using Mann-Whitney U test (*p < 0.05, **p < 0.01).

The findings implicate, but do not definitively demonstrate a role for LOX and PLOD2 in generating the increased level and greater complexity of collagen crosslinking we quantified in the more aggressive human breast tumors. Nevertheless, the combination of these gene expression data with our findings showing increased collagen crosslink abundance provide compelling evidence to suggest that increased expression of LOX and PLOD2, particularly from stromal cells, likely contribute to elevated levels of collagen crosslinks and HLCCs in TN tumors. Moreover, given that TN tumors are the most aggressive and lethal breast cancer, these data link human breast tumor aggression to increased levels of total and complex collagen crosslinks and higher stromal stiffness. Accordingly, the findings implicate the collagen crosslinking enzymes LOX and LH2, and by extension factors that regulate their expression, in breast cancer aggression.

### Stromal — and not epithelial — crosslinking enzymes regulate tissue fibrosis and collagen crosslinking *in vivo*

Prior studies demonstrated that both cancer cell lines and stromal fibroblasts express LOX and LH2 to induce tissue stiffening and fibrosis implying they also drove collagen cross-linking. Seminal articles identified hypoxia-induced HIF1a as a key regulator of tumor epithelial LOX and LH2 expression and suggested epithelial secretion of these enzymes drives collagen remodeling, crosslinking and stiffening that foster tumor cell dissemination and primes the pre-metastatic niche to facilitate metastatic colonization (22,24,25). Our analysis of breast cancer clinical specimens showed that the stromal cells in the breast tissue express higher levels of LOX and PLOD2 as compared to the breast tumor epithelium. Moreover, our prior studies showed LOX is expressed in stromal fibroblasts in a transgenic mouse model of ErbB2-induced mammary tumor malignancy. Furthermore, we demonstrated that fibroblasts expressing LOX injected into a cleared mammary fat pad not only induced ECM remodeling and stiffening but also potentiated the growth and malignant progression of pre-malignant tumor cells injected into the modified glands (12). Thus, the relative contribution of tumor- and stromal-derived collagen crosslinking enzymes to tumor fibrosis, ECM remodeling and collagen crosslinking remains unclear; particularly in the context of spontaneous tumors and patient tumors in which the ECM evolves concurrently with tumor progression.

To directly test the extent to which epithelial-derived LOX can crosslink collagen and induce tissue fibrosis and stromal stiffening in mammary tissue, we created a genetically engineered mouse model (GEMM) in which we targeted and controlled luminal epithelial-specific expression of mouse LOX using the MMTV-rtTA promoter (Epithelial LOX overexpression [OX]). (**Fig. 3a, Suppl. Fig. 6a-b**). We crossed these mice into the PyMT spontaneous mammary tumor model to enhanced LOX expression in the mammary tumor epithelium (PyMT epithelial LOX OX) and assayed their stromal phenotype as compared to the mammary glands from age-matched PyMT control mice (PyMT). Again, despite confirming ectopic LOX expression and elevated levels of cleaved LOX protein in the mammary tumor epithelial compartment (**Suppl. Fig 6c**), we were not able to detect any increase in the levels, nor any altered organization of the mammary gland interstitial collagen (**Fig 3b-c**). Moreover, xAAA crosslinking analysis revealed that the level of collagen crosslinks between the PyMT control and PyMT epithelial LOX OX mammary gland stroma were indistinguishable (**Fig. 3d-i**). Nevertheless and importantly, we could easily and consistently detect a significant increase in fibrillar collagens, collagen crosslinks and stromal stiffness in the PyMT mammary tumors as compared to age-matched FVB mammary glands lacking tumors (**Fig. 4c,f-k).** These studies both validate the sensitivity of our crosslinking assay and imply that tumor epithelial LOX is not the primary driver of collagen crosslinking and stiffening in endogenous mammary tumors.

**Figure 3:**
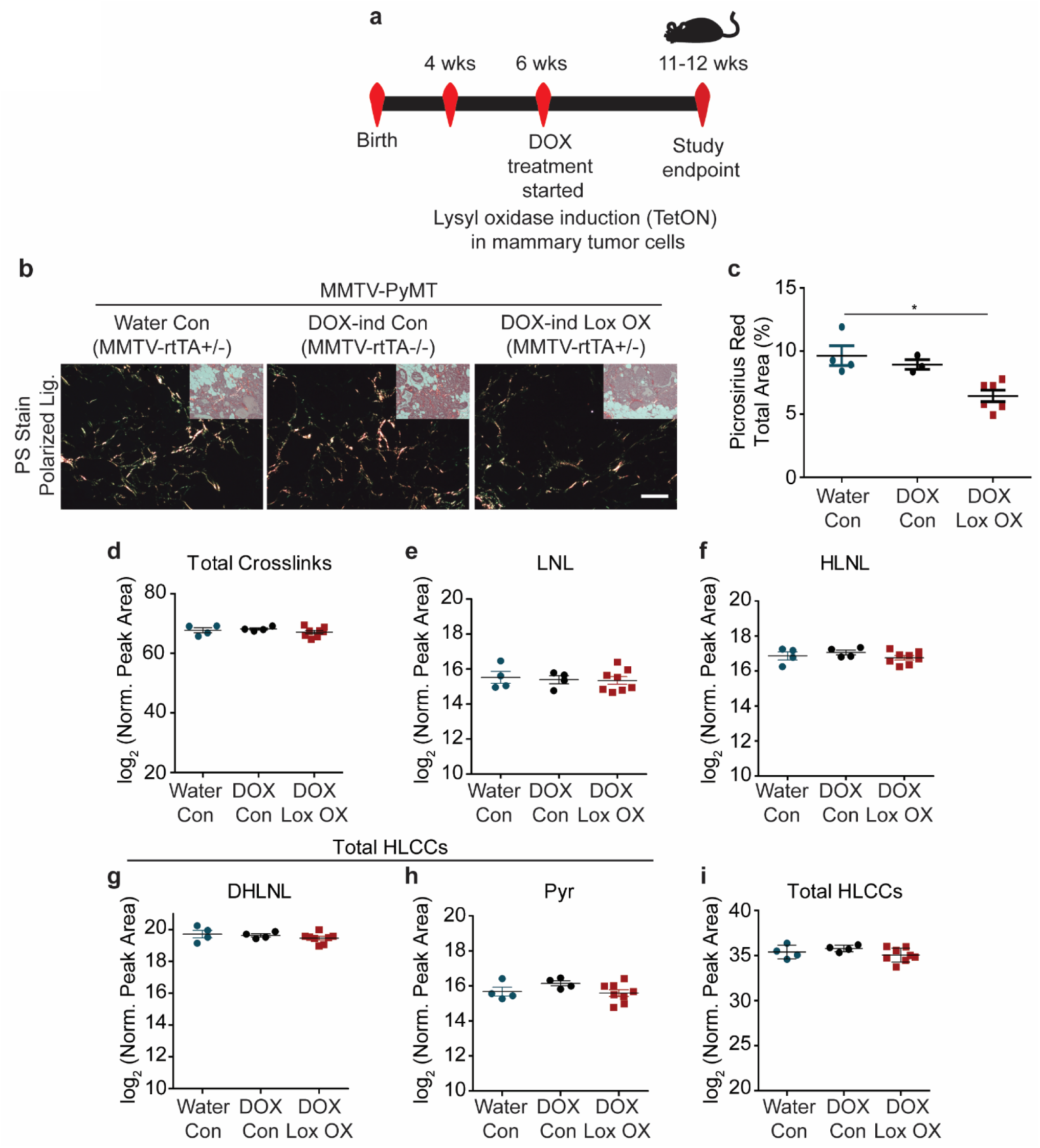
Epithelial-derived collagen crosslinking enzymes fail to induce collagen crosslinking. (**a**) Schematic depicting the experimental strategy used to induce epithelial Lox overexpression. (**b**) Polarized light images with brightfield inset of picrosirius red stained murine mammary tissues. (**c**) Quantification of percent area of picrosirius red staining per field of view, plotted as a scatter plot of the mean for each animal ± SEM. Statistical analysis was performed using Kruskal-Wallis one-way ANOVA (*p < 0.05). (**d**) Scatter plot showing individual and mean values ± SEM of total tissue collagen crosslinks in PyMT controls (Water n = 4, DOX n = 4) and PyMT epithelial Lox overexpression (n = 8). (**e-h**) Scatter plots showing individual and mean values ± SEM for each LCC and HLCC collagen crosslink measured in PyMT control and PyMT epithelial Lox overexpression tumor tissue. (**i**) Scatter plot showing individual and mean values ± SEM of total HLCCs calculated as the sum of DHLNL and Pyr crosslinks. Quantity of crosslinks per tissue was calculated normalizing crosslinks to total collagen content (i.e., hydroxyproline abundance) and wet tissue weight. Values were plotted as log_2_ transformed normalized peak areas as quantified from LC-MS data. Statistical analyses for crosslinking data were performed using one-way ANOVA for overall comparison and unpaired t-test for individual comparisons (*p < 0.05, **p < 0.01, ***p < 0.001, ****p < 0.0001).

**Figure 4:**
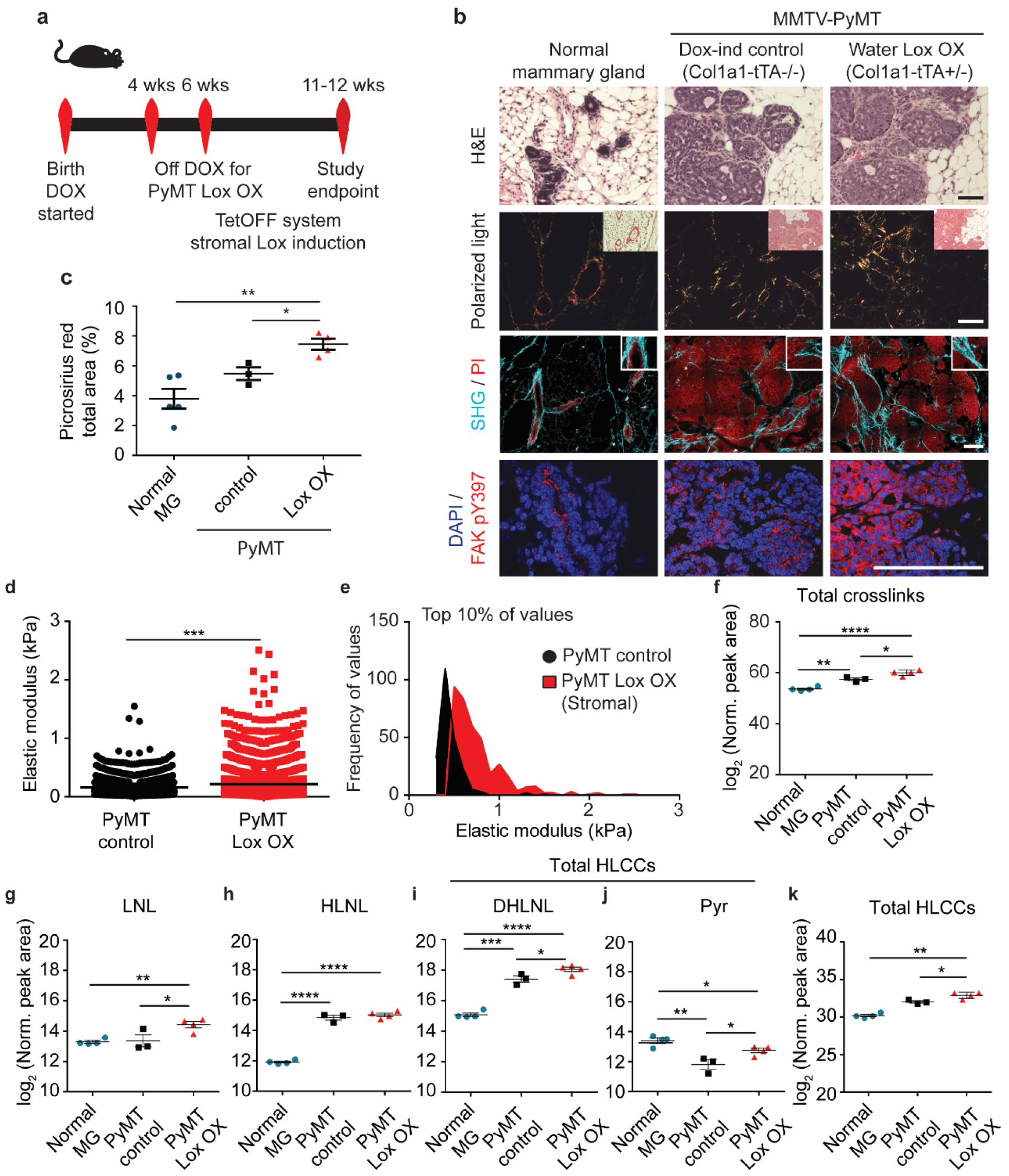
Stromal-derived LOX regulates collagen crosslinking and stiffening. (**a**) Schematic depicting the experimental strategy used to induce stromal Lox overexpression. (**b**) Representative images of normal murine mammary gland and PyMT control and stromal Lox overexpressing tumor tissues. (**Top row**) Brightfield images of H&E stained murine mammary tissues. (**Second row**) Polarized light images with brightfield inset of picrosirius red stained murine mammary tissues. (**Third row**) Two photon second harmonic generation (SHG) images of murine mammary tissues revealing collagen organization (turquoise) and propidium iodide (PI; red) stained nuclei. (**Bottom row**) Confocal images of mammary tissue stained with anti-FAK pY397 monoclonal antibody (red) and DAPI (blue; nuclei). Scale bars are all 100 µm. (**c**) Quantification of fibrillar collagen by picrosirius red staining by percent area per field of view. The mean of 3-5 regions was calculated and plotted for each animal ± SEM. Statistical analysis was performed using Kruskall-Wallis one-way ANOVA (*p < 0.05) for overall relationship and unpaired t-test for comparing individual groups (*p<0.05, **p<0.005). (**d**) Scatter plot showing individual values and mean of the top 10% of elastic modulus measurements performed by AFM microindentation on the mammary stroma from mice with PyMT-induced tumors (PyMT Control; n = 3 mice) as compared to the stroma in PyMT-induced tumors in which lysyl oxidase was elevated in stromal cells (PyMT Lox OX; n = 4 mice). Values reflect measurements taken from 3-5 individual force map regions per mammary gland. Statistical analyses were performed using Mann-Whitney U test (***p < 0.001). (**e**) Histogram showing the distribution of the top 10% of elastic modulus measurements by AFM microindentation in PyMT control and Lox OX tumors. Statistical analysis was performed using Mann-Whitney U test (****p < 0.0001). (**f**) Scatter plots showing individual and mean values ± SEM of total collagen crosslink abundance in the normal murine mammary gland (n = 4) as compared to glands with doxycycline-induced PyMT tumors (n = 3 mice per control group) and PyMT tumors in which stromal lysyl oxidase was elevated (n = 4 mice per Lox OX group). (**g-j**) Scatter plots showing individual and mean values ± SEM of LCC and HLCC crosslinks quantified in normal mammary gland, PyMT Control tumors and PyMT Lox OX tumors. (**k**) Scatter plot showing individual and mean values ± SEM of total HLCCs calculated as the sum of DHLNL and Pyr crosslinks. Quantity of crosslinks per tissue was calculated normalizing crosslinks to total collagen content (i.e., hydroxyproline abundance) and wet tissue weight. Values were plotted as log_2_ transformed normalized peak areas as quantified from LC-MS data. Statistical analyses for crosslinking data were performed using one-way ANOVA for overall comparison and unpaired t-test for individual comparisons (*p < 0.05, **p < 0.01, ***p < 0.001, ****p < 0.0001).

We and others have implicated stromal fibroblast LOX as a key promoter of epithelial tumor progression and aggression(12,14,40). To directly test the functionality of the ectopically-expressed LOX and the relevance of fibroblast-specific expression of LOX, we next created mouse cohorts of PyMT GEMMs in which we restricted ectopic LOX expression to the stromal population using the Col1a1-tTA promoter (MMTV-PyMT+/-; Col1a1-tTA+/-;TetO_mLox+/-; herein denoted PyMT LOX OX) (**Fig. 4a, Suppl. Fig. 7b**). To begin with, induction of LOX in the stromal cells markedly enhanced the amount of fibrillar collagen in the MMTV-PyMT LOX OX mammary glands, as revealed by quantification of polarized images of Picrosirius Red stained tissue (**Fig. 4b, second row images; quantified in c**). We also detected more and thicker linearized interstitial collagen when LOX was increased in the stromal cells by two-photon second harmonic generation imaging (**Fig. 4b, third row panels**). In addition, immunostaining revealed more phosphorylated tyrosine 397 focal adhesion kinase protein (^pY397^FAK) in the mammary epithelium of the glands in which stromal LOX was elevated (**Fig. 4b, bottom panels**), likely reflecting the increase in elasticity that we measured in the tissue stroma using atomic force microscopy (AFM) indentation(12) (**Fig. 4d-e & Suppl. Fig. 9b**). Consistently, we measured higher levels of total collagen crosslinks in the PyMT LOX OX mice as compared to the levels quantified in the MMTV-PyMT control glands (**Fig. 4f**). Furthermore, the most significant increases that we quantified in the PyMT LOX OX glands were dihydroxy lysinonorleucine (DHLNL) and pyridinoline (Pyr) crosslinks (**Fig. 4g-k**), which are the crosslinks generated through the HLCC pathway that promote mechanical stability and strength in skeletal tissue(37, 41). Of note, we did not detect any increase in DHLNL or Pyr in the mammary glands in which ectopic LOX expression was elevated in the mammary epithelium using the MMTV promoter (**Fig. 3g-h**). These findings imply that stromal cells are the primary regulators of interstitial collagen crosslinking and stromal stiffening in mammary tumors.

### Tissue inflammation regulates tissue fibrosis, collagen crosslinking and stromal stiffening

Cancer progression is accompanied by tissue inflammation and the most aggressive human breast tumors with the stiffest invasive stroma harbor the highest number of macrophages(2, 35). Consistently, decreasing the number of tumor macrophages, either through genetic ablation of macrophage colony stimulating factor (CSF-1) or via pharmacological and inhibitory antibody treatment with anti-CSF1 antibody, reduces lung metastasis in the PyMT mouse model of mammary cancer(33,34,42,43). Interestingly, macrophage ablation was more effective at preventing lung metastasis in the PyMT mammary tumor model when the treatment was initiated early, prior to malignant transformation and coincident with the onset of tissue fibrosis(33). These findings raise the intriguing possibility that macrophage ablation may regulate tumor aggression, at least in part, by promoting collagen remodeling and inducing ECM crosslinking and stromal stiffening (**Fig. 3 & 4**).

To assess the possibility that there is a causal association between macrophage-mediated tissue inflammation, tumor fibrosis and ECM stiffening and mammary tumor aggression, PyMT mice were treated with anti-CSF-1 antibody or a non-specific IgG control antibody commencing at four weeks of age, prior to the onset of ductal hyperplasia(44). Mouse cohorts (6/treatment group/time point) were sacrificed at eight and eleven weeks of age. Immunostaining confirmed efficient reduction of mammary tumor tissue macrophages at eight weeks of age, as evidenced by significantly reduced F4-80 immunostaining that was also evident in the eleven week old treated tissue (**Fig. 5a top panel, Suppl. Fig. 8a**). The excised lungs from the eleven week mice confirmed reduced frequency of lung metastasis in the anti-CSF-1 antibody-treated group (**Fig. 5b**), consistent with our prior work documenting a significant inhibition of lung metastasis in fourteen week old mice when anti-CSF-1 treatment was initiated at four weeks(33). The mammary glands from the eight and eleven week old mice were excised and analyzed for fibrosis and biomechanical properties (**Fig. 5a,c-i**). Despite confirming an equivalent number of fibroblasts in the treated and nontreated groups, polarized images of PS stained tissue revealed lower levels of total fibrillar collagen in the stroma of the eight week anti-CSF1 antibody treated group (**Fig. 5a, third row panel, Suppl. Fig. 10b-c**). AFM microindentation additionally demonstrated that the tissue stroma from both the eight week old (**Fig. 5d**), and the eleven week old (not shown) mammary glands was softer, likely accounting for the reduced integrin mechanosignaling detected in the CSF1-antibody treated tissue, as revealed by less intense staining for ^pY397^FAK **(Fig. 5a, second row panel**). Consistently, we also observed reduced levels of Lox mRNA in eight week, anti-CSF1 treated mice by in situ hybridization (**Fig. 5a, bottom panel, Fig. 5e**). In agreement with prior studies, nearly all the detected Lox mRNA was restricted to stromal cells (**Fig 5a, bottom panel**)(14). These findings suggest that reducing the level of tumor-associated macrophages (TAMs) not only prevents lung metastasis but also concomitantly reduces tissue fibrosis and stiffening, likely by preventing stromal cell activation.

**Figure 5:**
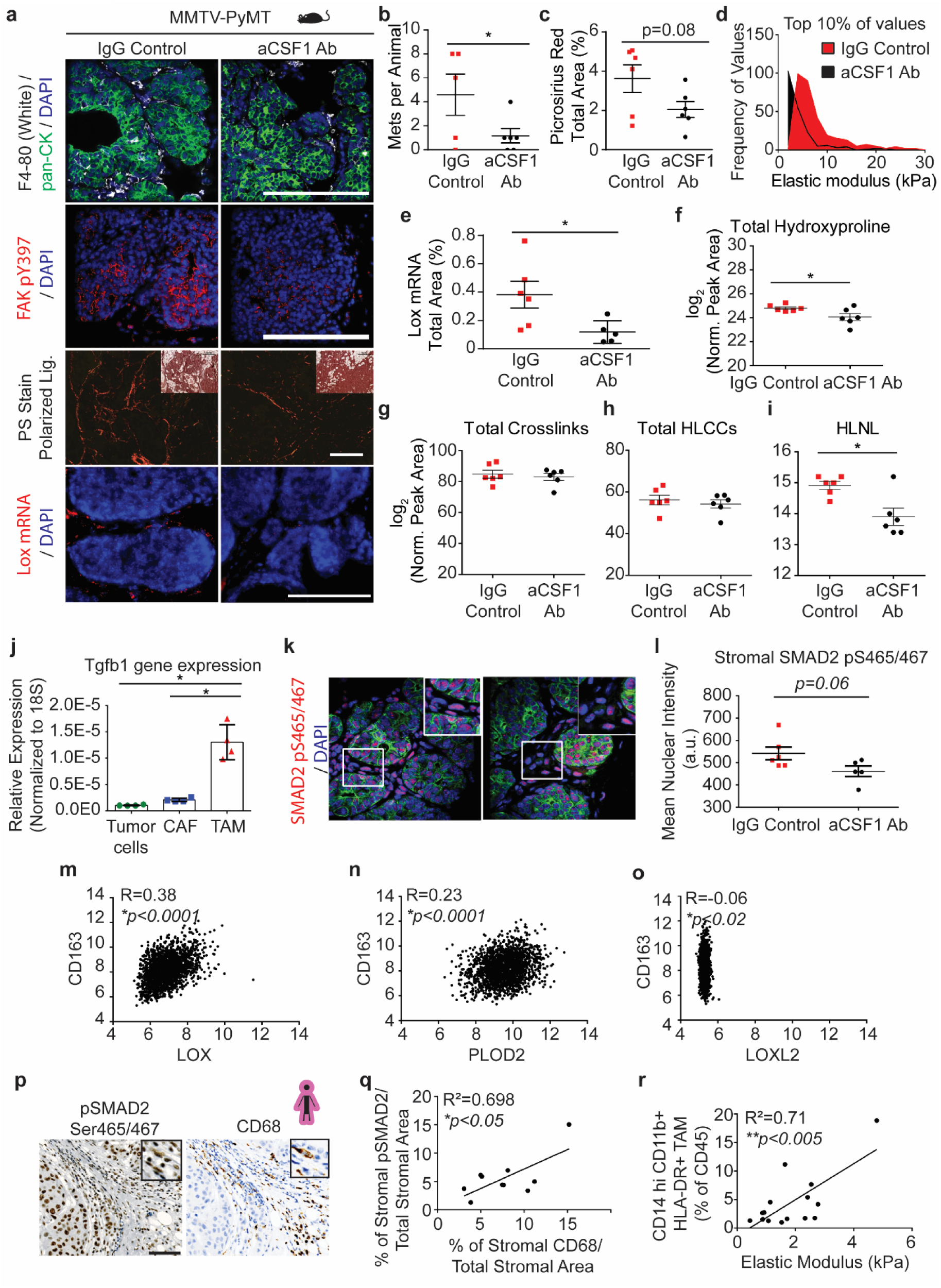
Tumor infiltrating macrophages secrete TGFb to activate stromal-mediated collagen crosslinking. (**a**) Representative images of PyMT tumor tissue from 8 weeks of age mice treated with anti-CSF1 blocking antibody or IgG1 control. (**Top row**) IgG1 treated (n = 6) and anti-CSF1 treated (n = 5) PyMT tumor tissue stained for pan-cytokeratin (green) marking epithelial cells, F4/80 (white) marking tumor infiltrating macrophages, and DAPI marking nuclei (blue). (**Second row**) IgG1 treated (n = 6) and anti-CSF1 treated (n = 5) PyMT tumor tissue stained for tyrosine 397 phosphorylated focal adhesion kinase (red) indicative of mechanosignaling and DAPI marking nuclei (blue). (**Third row**) Polarized light images with brightfield inset of IgG1 treated (n = 6) and anti-CSF1 treated (n = 6) PyMT tumor tissue stained with picrosirius red to visualize fibrillar collagen. (**Bottom row**) Lox mRNA in situ hybridization in IgG1 treated (n = 6) and anti-CSF1 treated (n = 5) PyMT tumor tissue and DAPI marking nuclei (blue). Scale bar for all images is 100µm. (**b**) Quantification of the number of metastatic colonies in the lung tissues from IgG1 treated (n = 5) and anti-CSF1 treated (n = 6) mice at 11 weeks of age via PyMT IHC assessing 5 layers (5 micron section; 5 sections per layer; 50-100 microns steps). Statistical analysis was performed using unpaired t-test (*p < 0.05). (**c**) Quantification of fibrillar collagen by picrosirius red staining by percent area per field of view in 8 week old mice treated with anti-CSF1 blocking antibody or IgG1 control. The mean of 3-4 regions was calculated and plotted for each animal ± SEM. Statistical analysis was performed using unpaired t-test (p = 0.08). (**d**) Histogram showing the distribution of the top 10% of elastic modulus measurements by AFM microindentation in PyMT IgG1 treated (n =6) and anti-CSF1 treated (n = 4) tumors. Statistical analysis was performed using Mann-Whitney U test (****p < 0.0001). (**e**) Quantification of Lox mRNA signal by percent area of signal per field of view in 8 week old mice treated with anti-CSF1 blocking antibody (n = 5) or IgG1 control (n = 6). The mean of 5-6 regions was calculated and plotted for each animal ± SEM. Statistical analysis was performed using an unpaired t-test (*p < 0.05). (**f-i**) Scatter plots showing individual and mean values ± SEM of the levels of total hydroxyproline (collagen content) (**f**), total collagen crosslinks (**g**), HLCCs (**h**), and HLNL crosslinks (**i**) in 8 week old IgG1 treated and anti-CSF1 treated PyMT tumors. Quantity of crosslinks per tissue was calculated normalizing crosslinks to wet tissue weight. Values were plotted as log_2_ transformed normalized peak areas as quantified from LC-MS data. Statistical analysis was performed using unpaired t-test (*p < 0.05). (**j**) Quantification of Tgfb1 gene expression by RT-qPCR in tumor cells, cancer-associated fibroblasts, and macrophages sorted out from PyMT tumors (n = 4). Gene expression was normalized to 18S. Statistical analysis was performed using Kruskal-Wallis one-way ANOVA for overall comparison and Mann-Whitney U test for individual comparisons (*p < 0.05, ***p < 0.001). (**k**) Representative images of PyMT tumor tissue from mice treated with IgG1 (n = 6) and anti-CSF1 (n = 5) stained for pan-cytokeratin (green) marking epithelial cells, SMAD2 pS465/467 (red), and DAPI marking nuclei (blue). (**l**) Scatter plot showing individual and mean values ± SEM of the mean nuclear intensity of pSMAD^S465/467^ in stromal cells of IgG1 treated (n = 6) and anti-CSF1 treated (n = 5) PyMT mice. The mean for each animal was calculated from 4-7 regions within the tumor. Statistical analysis was performed using unpaired t-test (p = 0.06). (**m-o**) Scatter plot depicting the Spearman correlation of CD163 gene expression with LOX (**m**), PLOD2 (**n**), and LOXL2 (**o**) in human breast tumors (n = 1904). (**p**) Representative IHC images of serial human breast tumor sections stained for CD68 (**top**) and pSMAD2^S465/467^ (**bottom**) counterstained with hematoxylin to mark nuclei. (**q**) Scatter plot depicting the linear regression correlation of stromal pSMAD2 IHC staining with stromal CD68 IHC staining in human breast tumors (n = 10). (**r**) Scatter plot depicting the linear regression of CD14^+^ CD11b^+^ HLA-DR^+^ tumor associated macrophage infiltrate with tumor elastic modulus as measured by AFM microindentation in human breast tumors (n = 15).

TAMs secrete abundant TGFβ that stimulates fibroblast differentiation to a myofibroblast phenotype and induces expression of collagen crosslinking enzymes including LOX(14). PCR analysis of flow activated cell sorted (FACS) cells from the transformed mammary glands of PyMT mice (11 weeks) confirmed that the TAMs expressed by far the highest levels of TGFβ, as compared to neoplastic epithelium and the cancer-associated fibroblasts (**Fig. 5j**). These findings suggest that the tumor infiltrating macrophages could promote fibrosis and stromal stiffening through secreted TGFβ; a finding supported by a strong stromal pSMAD2 (SMAD2 pS465/467) in the IgG control antibody-treated PyMT mammary glands (**Fig. 5k-l**). To examine whether TAM recruitment could regulate collagen crosslinking in human tumors we queried a publicly available human breast tumor gene expression data set for TAM markers and collagen modifying enzymes. Consistent with a role for TAMs in stimulating expression of collagen modifying enzymes, gene expression of the TAM marker CD163 positively correlated with both LOX and PLOD2 but not LOXL2 in human breast tumors (n = 1904) (**Fig. 5m-o**). Moreover, co-staining of resected human breast tumors with the macrophage/monocyte marker CD68 and the downstream TGFβ signaling molecule pSMAD2 Ser465/467 revealed a significant positive correlation between pSMAD2 Ser465/467 and infiltrating tumor macrophages at the invasive front of human breast tumors (**Fig. 5p-q**). Furthermore, FACS analysis established a significant correlation between the infiltrating tumor associated macrophages, as demonstrated by CD14^hi^CD11b^+^ HLA-DR^+^ cell surface markers normalized to total CD45 infiltrating cells, and the elastic modulus of the invasive front of human invasive breast cancers (**Fig. 5r**). These findings suggest that tumor inflammation and macrophage secreted factors such as TGFβ could promote tissue fibrosis by enhancing fibroblast expression of the collagen crosslinking enzymes LOX and PLOD2, but not LOXL2, to induce collagen crosslinking and stromal stiffening.

### LH2 inhibition reduces lung metastasis and stromal LH2 predicts poor prognosis in breast cancer patients

Using the data set generated from the micro-dissected tumors, we next explored the clinical relevance of epithelial vs. stromal LOX and PLOD2 expression by assessing their relative contribution to overall survival in breast cancer patients. Surprisingly, neither stromal cell nor epithelial LOX predicted overall patient survival in this cohort (**Fig. 6a-b**). However, the findings clearly showed that overexpression of stromal PLOD2, but not epithelial PLOD2, significantly correlated with poor breast cancer patient prognosis (**Fig. 6c-d**). The data provide further evidence that HLCCs, formed primarily through the activities of collagen crosslinking enzymes expressed by stromal cells, promote breast tumor aggression that contributes to poorer overall survival. The findings also highlight the importance of the collagen crosslinking profile and its potential impact on stromal stiffness in tumor aggression.

**Figure 6:**
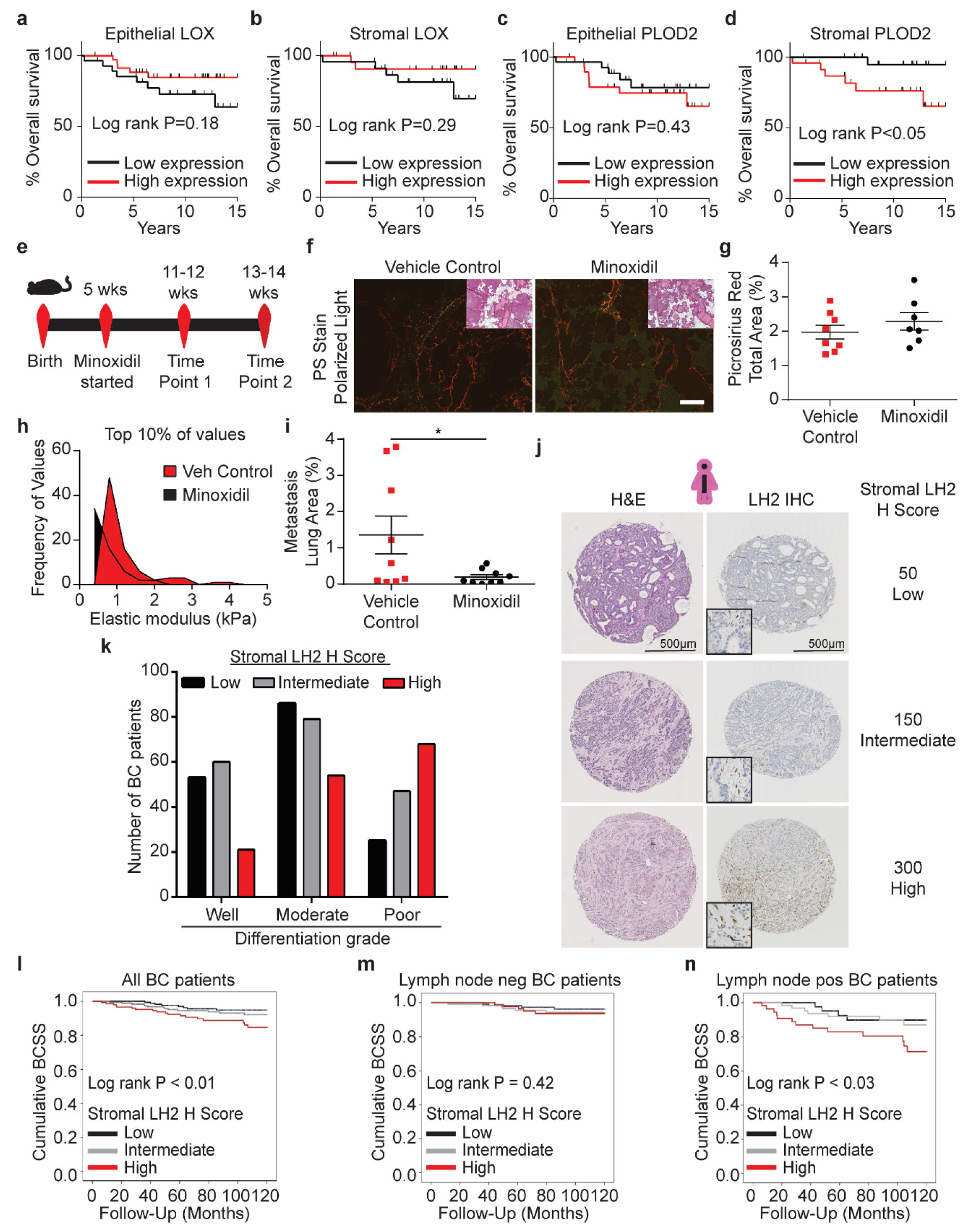
Stromal LH2 predicts poor patient outcomes. (**a-b**) Kaplan-Meier plots showing overall survival for patients based on levels of LOX expression in epithelial cells (low n = 28, high n = 36) (**a**) or stromal cells (low n = 23, high n = 24) (**b**). The median level of expression was defined as the cutoff for low and high expression. (**c-d**) Kaplan-Meier plots showing overall survival for patients based on levels PLOD2 expression in epithelial cells (low n = 28, high n = 29) (**c**) or stromal cells (low n = 23, high n = 24) (**d**). The median level of expression was defined as the cutoff for low and high expression. (**e**) Schematic depicting the experimental timeline used to inhibit lysyl hydroxylase 2 in PyMT mice. (**f**) Representative polarized light images with brightfield insets of picrosirius red stained tumor tissue from PBS vehicle treated control (n = 8) and minoxidil treated (n = 7) PyMT mice. Scale bar is 100um. (**g**) Quantification of fibrillar collagen by picrosirius red staining by percent area per field of view. The mean was calculated and plotted for each animal ± SEM. (**h**) Histogram showing the distribution of the top 10% of elastic modulus measurements by AFM microindentation in PyMT control and Lox OX tumors. Statistical analysis was performed using Mann-Whitney U test (****p < 0.0001). (**i**) Scatter plot quantifying the area of lung sections occupied by metastases from vehicle treated (n = 9) and minoxidil treated (n = 8) mice at 13 weeks of age via H&E staining and assessing 4 layers (5 micron section; 5 sections per layer; 50-100 microns steps). Statistical analysis was performed using a two-tailed unpaired t-test (*p < 0.05). (**j**) Representative phase contrast images of sections from tissue microarrays (TMAs) of human breast cancers representing incident breast cancer cases collected and arrayed as 1-mm cores from each tumor. Sections were stained with Hematoxylin and Eosin (H&E; top) and lysyl hydroxylase two (LH2; bottom) via immunohistochemistry. (**k**) Bar graphs showing clinical correlation between lysyl hydroxylase two (LH2) score as a function of tumor grade (*see Table 1 for number of patients*). LH2 IHC staining was assessed with the semi-quantitative stromal specific H-score from 0 to 300. The lowest tertile of LH2 H-scores was defined as H-scores between 0 and less or equal to 120, the intermediate H-score to above 120 and equal or less than 230, and the highest stromal LH2 score as above 230. For tumor grade and LH2 H score, statistical analysis was performed using a linear-by-linear association (\*\*\**P<0.0001*). (**l**) Kaplan-Meier curves indicating cumulative breast cancer specific survival (BCSS) based on stromal LH2 H score assessed in breast cancer patients up to 10 years after diagnosis (LH2 low n = 175, intermediate n = 188, high n = 146). (**m**) BCSS curves by stromal LH2 H score including only axillary lymph node negative patients (LH2 low n = 116, intermediate n = 116, high n = 90). (**n**) BCSS curves by stromal LH2 H score including only axillary lymph node positive patients (LH2 low n = 44, intermediate n =63, high n = 54). For Kaplan-Meier curves, statistical analyses were performed by LogRank test.

LH2 (the protein encoded by PLOD2) is a key enzyme that regulates the level of HLCC crosslinking of fibrillar collagen and consequently contributes substantially to the tensile properties of the tissue stroma(19,23,45). We quantified the highest level of HLCC crosslinking and the stiffest invasive front in the breast tissue from women with the most aggressive breast cancers (**Fig. 1j**). Consistently, when we analyzed the gene expression data from a published cohort of breast cancer patients we observed that ER-/HER2-(TN; n=133) breast cancers express the highest level of the PLOD2 gene transcript as compared to the levels expressed in HER2+ (n=73) and ER+/HER2-(luminal, n=314) breast tumors (**Suppl. Fig. 11a**)(46). Further analysis showed that high PLOD2 expression in breast cancer patients with HER2+ and TN tumors significantly predicts reduced distant metastasis-free survival (DMFS), as well as increased risk of relapse in TN tumors (**Suppl. Fig. 11b-d**)(46, 47). These findings implicate PLOD2/LH2 as a key regulator of breast tumor aggression through its ability to induce the HLCC crosslinking of fibrillar collagens that enhance the stiffness of the tumor stroma.

To evaluate whether LH2 does enhance stromal stiffness to promote breast tumor aggression, we systemically treated a cohort of PyMT mice with the LH2 inhibitor minoxidil or vehicle (PBS) from five weeks of age until sacrifice at either 11-12 weeks or 13-14 weeks of age (**Fig. 6e**). Consistent with its predicted role in enhancing stromal stiffness through modifying the nature of collagen crosslinks, AFM microindentation revealed that the phenotypically-similar collagen-rich ECM adjacent to tumors from the minoxidil treated mice (see polarized imaging of picrosirius red stained tissue) was significantly softer when compared to the vehicle-treated age/stage-matched PyMT mice (**Fig. 6f-h**). Furthermore, inhibiting LH2 also significantly decreased lung metastasis, causally linking HLCC collagen crosslinking to stiffness-mediated breast tumor aggression (**Fig. 6i**). Our gene expression data from the epithelial and stromal microdissection cohort indicated that stromal PLOD2, but not epithelial PLOD2, strongly predicted poor survival in breast cancer patients. Therefore, we next sought to definitively assess the relative contribution of stromal versus neoplastic epithelial LH2 protein expression to human breast tumor aggression using a large tissue array of annotated human breast tumor biopsies. To accomplish this, we first developed an LH2 immunostaining protocol (IHC) and then we analyzed neoplastic epithelial and stromal expression of the LH2 enzyme in tissue biopsies from a large cohort of histopathologically classified breast cancer patients (N=505) with accompanying clinical information and follow up data (**Suppl. Table 2** and **Suppl. Table 3**). LH2 IHC staining in the epithelium and stromal cells was scored as low, intermediate and high and the relationship between epithelial versus stromal expression and breast cancer patient outcome was calculated (**Fig. 6j; Suppl Fig. 12a**). IHC scoring analysis revealed that only a modest number of patients with poorly differentiated breast tumors expressed moderate to high levels of epithelial LH2 (**Suppl. Fig. 12b**); a finding that accords with prior links between tumor size, hypoxia and LH2-dependent tumor aggression(23, 24). More strikingly however, we observed that a disproportionately high number of breast cancer patients with moderately and poorly differentiated breast tumors had high stromal cell LH2 (indicated by an H score of above 230; **Fig. 6k**). These data and our observation that even well differentiated breast tumors express intermediate levels of stromal cell LH2 (H score above 120 and equal or less than 230) imply that stromal cell LH2, rather than epithelial LH2, may be a more robust indicator of breast tumor aggression. Consistently, stratification of breast tumor patient analysis into ER+/HER2-(N=296); HER2+ (N=36) and TN (N=32) showed an enrichment for intermediate and high stromal LH2 H-score in the more aggressive cancer subtypes (**Suppl. Table 2** and **Suppl. Table 3**). Furthermore, we uncovered a significant correlation between high stromal cell LH2 H-score and shorter breast cancer patient-specific survival when adjusted for age at diagnosis (**Fig. 6l)**. We also established a significant association between stromal cell but not epithelial LH2 levels and survival in lymph node positive breast cancer patients but not lymph node negative breast cancers (**Fig. 6m-n; Suppl Fig. 12d-e**). These clinical data identify stromal cell LH2 as a novel biomarker with potential to predict metastatic disease and poor patient survival among breast cancers overall, as well as within the highest risk TN breast cancer subtype.

## Discussion

We identified infiltrating macrophages as key regulators of stromal cell-mediated collagen crosslinking, stromal stiffening and tumor metastasis. Our clinical findings corroborated our experimental results, revealing significant associations between macrophages and collagen crosslinking enzymes providing evidence for a critical impact of inflammatory stromal cell-mediated collagen crosslinking and stromal stiffening in tumor aggression and patient outcome. Diverse subsets of myeloid cells account for adverse patient outcomes because they differentially promote angiogenesis, tumor cell intravasation, and suppress the anti-tumor immune response(48). Our results expand this perspective to include a key role for early infiltrating macrophages in initiating a collagen crosslinking and stiffening program that ultimately fosters tumor aggression and progression. We also determined that TN breast cancers contain much higher amounts of LH2- and LOX-derived collagen crosslinks. Specifically, we demonstrated that significantly higher levels of HLCC crosslinks could explain their higher stromal stiffness and aggressiveness(2). Intriguingly, in depth collagen analysis suggested that each breast tumor subtype exhibits a distinct collagen organization, stiffness and crosslinking profile, raising the possibility that distinct collagen architectures and crosslinking signatures may reflect differences in tissue pathobiology. Indeed, TN breast cancers often present with high macrophage infiltration; an observation that is consistent with our AFM, two photon and collagen crosslinking analysis data(2). Given that the immune cell infiltrate significantly influences treatment response it is tempting to speculate that distinct collagen architectures, crosslinking signatures and stiffnesses similarly regulate therapeutic efficacy.

We determined that stromal LH2 is a robust predictor of survival in breast cancer patients, especially in those that are lymph node positive, supporting a potentially important clinical link to stromal collagen crosslinking. Our data also revealed that both the upregulation of crosslinking enzymes and in turn, collagen crosslinking, occur at an early stage of malignancy that is concurrent with tumor cell invasion and coincident with the accumulation of infiltrating macrophages(49). In part, this may explain why therapeutics targeting LOX and LOXL2 have thus far failed to significantly prolong cancer patient survival. Substantial levels of collagen crosslinking are likely to have occurred prior to therapy administration, and while these agents may prevent further crosslinking activity, they are not capable of reversing crosslinks or collagen modifications already present in the tissue. The early initiation of collagen crosslinking also suggests that a detectable increase in stromal LH2 may provide an early prognostic marker of disease progression and aggression that could inform treatment strategies. To that end, our results show for the first time that LH2-derived collagen crosslinks are not only a distinct feature of enhanced stromal stiffening in TN breast cancer but are able to predict distant metastasis-free survival. These results also suggest that the profile of collagen crosslinks, and not simply crosslink abundance alone, may play a role in promoting tumor aggression, thus warranting further investigations into the contributions of HLCCs to ECM mechanical properties and tumor aggression.

Our findings identify fibroblasts as the dominant cell population that promotes reorganization and crosslinking of interstitial collagen to stiffen the tissue stroma. Only stromal-targeted, and not epithelial-targeted, tissue-specific inducible LOX produced any measurable change in collagen architecture, stiffness or crosslink abundance in a spontaneous tumor (25,50,51). Our conclusion is consistent with earlier work in which we failed to detect any alterations in fibrosis, collagen organization or stiffness when LOXL2 was either genetically-ablated, or ectopically-increased in the mammary epithelium of PyMT mouse tumors, despite documenting a significant impact on metastasis(40). Nevertheless, it is possible that our bulk assay would not detect any small increase in collagen crosslinking induced by invading tumor cells expressing high levels of LOX. To this end, immortalized tumor cells engineered to overexpress LOXL2 injected orthotopically as a bolus of cells were able to induce fibrosis and quantifiable changes in collagen remodeling and tissue stiffening; a finding we too confirmed using isolated PyMT tumor cells overexpressing the LOXL2 enzyme(40). Nevertheless, our data support the notion that in spontaneous tumors, the proximity of stromal cells to collagen and their significantly higher expression of crosslinking enzymes dictate the profile and extent of collagen crosslinking. Given that LH2 modifies collagen intracellularly and stromal cells secrete the vast majority of the interstitial fibrillar collagen, it is not surprising that this is the case(52, 53). Moreover, our results stress the necessity of choosing an appropriate model to study tumor associated ECM remodeling and suggest that orthotopic models may fail to accurately recapitulate the natural evolution of ECM in tumor progression.

Our findings not only underscore the importance of stromal stiffness and collagen crosslinking in cancer-associated fibrosis and disease aggression, but also define in more detail the molecular nature of collagen modifications that accompany pathological fibrosis. Clarifying the nature of collagen modifications and specific mechanisms regulating those alterations in fibrosis will assist in the strategic design of novel, efficacious strategies to combat progressive fibrosis and should prove instrumental in enabling further experimentation to understand pathological fibrosis. Indeed, defining the molecular regulators that stimulate collagen cross-linking, and mechanisms that distinguish resolvable from non-resolvable fibrosis would identify attractive therapeutic targets for several pathological fibrotic diseases with limited treatment options.

## Methods

### Human breast specimen acquisition and processing

Fresh human breast specimens from breast reduction, prophylaxis, or breast tumor mastectomy were either embedded in an optimum cutting temperature (OCT) aqueous embedding compound (Tissue-Plus, Scigen, Cat# 4583) within a disposable plastic base mold (Fisher, Cat# 22-363-554) and were snap frozen by direct immersion into liquid nitrogen and kept at −80°C freezer until cryo-sectioning for analysis, or specimens were formalin fixed and paraffin embedded (FFPE). All human breast specimens were collected from prospective patients undergoing surgical resection at UCSF or Duke University Medical Center between 2010 and 2014. The selected specimens were de-identified, stored, and analyzed according to the procedures described in Institutional Review Board (IRB) Protocol #10-03832 and #10-05046, approved by the UCSF Committee of Human Resources and the Duke’s IRB (Pro00034242)(2).

### Mouse studies

Macrophages were depleted in MMTV-PyMT mice by i.p. injections of 1mg of anti-CSF1 antibody clone 5A1 or an IgG1 control every 7 days starting at 4 weeks of age. Mice were sacrificed at 8 and 11 weeks of age for tissue analysis.

Minoxidil and PBS vehicle was administered by i.p. injections of 3 mg/kg minoxidil in PBS three times per week starting at 6 weeks of age. Mice were sacrificed at 11-12 and 13-14 weeks of age for tissue analysis.

### Generation of mice

All mouse studies were maintained under pathogen-free conditions and performed in accordance with the Institutional Animal Care and Use Committee and the Laboratory Animal Research Center at the University of California, San Francisco.

### TetO-mLOX-eGFP construct and transgenic mouse generation

Full length mouse Lox (mLox) cDNA was purchased from OriGene. The full length ORF was amplified by PCR using forward and reverse primers respectively: GCAGGGATCCGCCACCATGCGTTTCGCCTGGGCTG and GGCGTCTAGAGCACCATGCGTTTCGCCTGGGCTGTGC. Following digestion with BamHI and XbaI, the PCR product was inserted into pSK TetO IRES 3xnlsEGFP(54) downstream of the Tet regulated minimal CMV promoter and a 5’ UTR containing a chimeric intron of human β-globin and immunoglobulin heavy chain genes, which was expressed as a bicistronic mRNA via an internal ribosome entry site (IRES2) with eGFP targeted to the nucleus by a N terminal in frame fusion of 3 tandem repeats of the SV40 nuclear localization sequence (nls). The fragment containing the expression cassette from Tet regulated promoter to SV40 polyadenylation signal (SV40pA) was agarose gel purified from XhoI– EagI digested donor plasmid and was used to generate TetO-mLox-eGFP transgenic mice by pronuclear injection into FVB/n oocyte (Mouse Biology Program at UC Davis; project number MBP-834; colony number PN663).

### Generation of MMTV-PyMT/Col1a1(2.3)-tTA/TetO-mLox-eGFP mice

The MMTV-PyMT/Col1a1(2.3)-tTA/TetO-Rs1 triple transgenic mice were generated by heterozygote or homozygote crosses of mice carrying the TetO-mLox-eGFP transgene with mice carrying heterozygote of the Col1a1(2.3)-tTA transgene (line 139)(55) or MMTV-rtTA and MMTV-PyMT transgene(56) to generate the experimental triple-transgenic genotype. In all breeding thereafter, MMTV-PyMT/Col11a1-tTA or MMTV-rtTA/TetO-mLox-eGFP male mice were crossed with TetO-mLox-eGFP female mice. Two mg/mL doxycycline hyclate (Alfa Aesar; Cat# J60579) was added to 5% sucrose water to modulate TetO-mLox-eGFP transgene expression.

### Preparation of human breast specimens for hydrolysis

OCT was removed from tissue blocks by first transferring biospecimens to a conical tube and then performing 5X washes with 70% ethanol followed by 5X washes with 18 MΩ H_2_O^48^. Each wash consisted of vortexing the sample for 15 minutes at 4°C and then centrifuging at 18,000 x g for 15 minutes at 4°C. Between 1 and 3 milligrams of tissue was washed with 1X PBS buffer by vortexing for 15 minutes at 4°C and then sonicated on ice for 20 seconds using a Sonic Dismembrator M100 (ThermoFisher, San Jose, CA, USA). The homogenate was then centrifuged at 18,000 x g for 20 minutes at 4°C. The supernatant was removed and the pellet was re-suspended in 1mg/mL NaBH_4_ (prepared in 0.1N NaOH) in 1X PBS for 1 hour at 4°C with vortexing. The reaction was neutralized by adding glacial acetic acid to a final concentration of 0.1% (pH ∼ 3 −4)(57). The sample was then centrifuged at 18,000 x g for 20 minutes at 4°C. The supernatant was removed and the pellet was washed three times with 18 MΩ H_2_O to remove residual salt that could interfere with downstream LC-MS/MS analysis. The remaining pellet was dried under vacuum for further analysis.

### Protein hydrolysis

The dried sample was placed in a glass hydrolysis vessel and hydrolyzed in 6N HCl, 0.1% phenol. The hydrolysis vessel is flushed with N2 gas, sealed and placed in a 110°C oven for 24 hours. After hydrolysis, the sample was cooled to room temperature and then placed at −80°C for 30 minutes prior to lyophilization. The dried sample was re-hydrated in 100μL of 18 MΩ H_2_O for 5 minutes, then 100 μL of glacial acetic acid for 5 minutes and finally 400 μL of butan-1-ol for 5 minutes. Importantly, 10 μL of sample is removed after re-hydration in water and saved for determination of hydroxyproline content.

### Preparation of crosslink enrichment column

CF-11 cellulose powder is loaded in a slurry of butan-1-ol: glacial acetic acid, water (4:1:1) solution onto a Nanosep MF GHP 0.45μm spin columns until a settled resin bed volume of approximately 5mm is achieved. The resin is washed with 1.5 mL 4:1:1 organic mixture using an in-house vacuum manifold set up. Re-hydrated samples are then loaded onto individual columns, the vacuum is turned on and the sample is pulled through the resin into glass collection vials. The flow through is again passed over the resin to ensure maximal binding of crosslinked amino acids and set aside. The column is then washed with 1.5 mL of fresh 4:1:1 organic mixture. A fresh collection vessel is placed under the column and 750 μL of 18 MΩ H2O is used to elute crosslinked amino acids off of the CF-11 resin. The eluent is then placed in a speed vac and run until complete dryness. Dried eluent is then reconstituted in a buffer appropriate for downstream MS analysis on amide HILIC UHPLC columns.

### UHPLC analysis

Up to 20 μL of tissue hydrolysates were analyzed on a Vanquish UPHLC system (ThermoFisher, San Jose, CA, USA) using an Acquity UHPLC BEH Amide column (2.1 x 100mm, 1.7 μm particle size – Waters, Milford, MA, USA). Samples were separated using a 5 minute gradient elution (55% - 40% Mobile phase B) at 250 μL/min (mobile phase: (A) 10mM ammonium acetate adjusted to pH 10.2 with NH_4_OH (B) 95% acetonitrile, 5% Mobile Phase A, pH 10.2, column temperature: 35°C.

### MS data acquisition

The Vanquish UPHLC system (ThermoFisher, San Jose, CA, USA) was coupled online with a QExactive mass spectrometer (Thermo, San Jose, CA, USA), and operated in two different modes – 1. Full MS mode (2 μscans) at 70,000 resolution from 75 to 600 m/z operated in positive ion mode and 2. PRM mode at 17,500 resolution with an inclusion list of in-tact crosslinked amino acid masses (Supplementary Table 2), and an isolation window of 4 m/z. Both modes were operated with 4 kV spray voltage, 15 sheath gas and 5 auxiliary gas. Calibration was performed before each analysis using a positive calibration mix (Piercenet – Thermo Fisher, Rockford, IL, USA). Limits of detection (LOD) were characterized by determining the smallest injected crosslinked amino acids (LNL, DHLNL, d-Pyr,) amount required to provide a signal to noise (S/N) ratio greater than three using < 5 ppm error on the accurate intact mass. Based on a conservative definition for Limit of Quantification (LOQ), these values were calculated to be threefold higher than determined LODs.

### MS data analysis

MS Data acquired from the QExactive were converted from a raw file format to .mzXML format using MassMatrix (Cleveland, OH, USA). Assignment of crosslinked amino acids was performed using MAVEN (Princeton, NJ, USA)(58). The MAVEN software platform provides the means to evaluate data acquired in Full MS and PRM modes and the import of in-house curated peak lists for rapid validation of features. Normalization of crosslinked amino acid peak areas was performed using two parameters, 1. Hydroxy proline content and 2. Tissue dry weight pre-hydrolysis (in milligrams)(57). Hydroxy proline content is determined by running a 1:10 dilution of the pre-enrichment sample through the Full MS mode (only) described above and exporting peak areas for each run.

### Quantification of crosslinked amino acids

Relative quantification of crosslinked amino acids was performed by exporting peak areas from MAVEN into GraphPad (La Jolla, CA, USA) and normalizing based on the two parameters described above. Statistical analysis, including T test and ANOVA (significance threshold for P values <0.05) were performed on normalized peak areas. Total crosslink plots were generated by summing normalized peak areas for all crosslinks in a given sample. Total HLCC plots were generated by summing normalized peak areas for all HLCC (DHLNL, Pyr, dPyr) crosslinks in a given sample.

### Picrosirius red staining and quantification

FFPE tissue sections were stained using 0.1% Picrosirius red (Direct Red 80, Sigma-Aldrich, Cat# 365548 and picric acid solution, Sigma-Aldrich, Cat# P6744) and counterstained with Weigert’s hematoxylin (Cancer Diagnostics, Cat# CM3951), as previously described(2). Polarized light images were acquired using an Olympus IX81 microscope fitted with an analyzer (U-ANT) and a polarizer (U-POT, Olympus) oriented parallel and orthogonal to each other. Images were quantified using an ImageJ macro to determine percentage area coverage per field of view. The ImageJ macro is available at https://github.com/northcottj/picrosirius-red.

### Second harmonic generation image acquisition

Second harmonic generation (SHG) imaging was performed using a custom-built two-photon microscope setup equipped resonant-scanning instruments based on published designs containing a five-PMT array (Hamamatsu, C7950), as previously published(2). The setup was used with two channel simultaneous video rate acquisition via two PMT detectors and an excitation laser (2W MaiTai Ti-Sapphire laser, 710–920 nm excitation range). SHG imaging was performed on a Prairie Technology Ultima System attached to an Olympus BX-51 fixed stage microscope equipped with a 25X (NA 1.05) water immersion objective. Paraformaldehyde-fixed or FFPE tissue sections were exposed to polarized laser light at a wavelength of 830 nm and emitted light was separated using a filter set (short pass filter, 720 nm; dichroic mirror, 495 nm; band pass filter, 475/40 nm). Images of x–y planes at a resolution of 0.656 mm per pixel were captured using at open-source Micro-Magellan software suite.

### Immunofluorescence/Immunohistochemistry

Immunofluorescence staining was performed as previously described(2). Briefly, mouse tissues were harvested and fixed with 10% buffered formalin phosphate (Fisher, Cat# 100-20) for 16-24 hours at room temperature and then further processed and paraffin-embedded. Five-µm sections dried for 30 minutes in 60^°^C, follow by deparaffinization and rehydration. Antigen retrieval was perform using DAKO Target Retrieval Solution (DAKO, Cat# S1699) for five minutes in a pressure cooker set to high pressure. Tissue sections were incubated with anti-FAK pY397 antibody (Abcam, Cat# Ab39967, dilution 1:25) overnight at 4°C and with anti-rabbit IgG Alex Fluor 633 (ThermoFisher, Cat# A-21070, dilution 1:2000) for one hour at room temperature. Antigen retrieval for immunofluorescent staining of SMAD2 pS465/467, cytokeratin 8+18, cytokeratin 5, F4/80, vimentin, and PDGFRα was performed using Diva Decloaker (BioCare, Cat# DV2004MX) for five minutes in a pressure cooker set to high pressure. Tissue sections were incubated with anti-SMAD2 pS465/467 antibody (Millipore, Cat# AB3849-I, dilution 1:100), anti-F4/80 antibody (AbD Serotec, clone CI:A3-1, Cat# MCA497GA, dilution 1:400), anti-cytokeratin 8+18 antibody (Fitzgerald, Cat# 20R-CP004, dilution 1:400), anti-cytokeratin 5 antibody (Fitzgerald, Cat# 20R-CP003, dilution 1:400), anti-vimentin antibody (Cell Signaling, Cat# 5741, dilution 1:100), and anti-PDGFRα (CD140a) antibody (Biolegend, Cat# 135901, dilution 1:100) overnight at 4°C and with anti-rat IgG Alex Fluor 488 (ThermoFisher, Cat# A-11006, dilution 1:1000), anti-guinea pig IgG Alex Fluor 568 (ThermoFisher, Cat# A-11075, dilution 1:1000), anti-rabbit IgG Alex Fluor 633 (ThermoFisher, Cat# A-21070, dilution 1:2000), anti-rabbit IgG Alex Fluor 633 (ThermoFisher, Cat# A-21070, dilution 1:1000) for one hour at room temperature.

Quantification of stromal nuclear SMAD2 pS465/467 was performed using Imaris 9. Surfaces were created around each nucleus and epithelial nuclei were manually excluded based on cytokeratin signal and cell morphology. The means of the mean nuclear signal intensity for all stromal nuclei were calculated for each field of view and averaged for every animal. Lungs from 11 week old IgG1 control and anti-CSF1 treated PyMT mice were cut into 5 micron sections from 5 layers with 100 microns between the first three layers and 50 microns between the last two layers. Sections were analyzed for metastases by PyMT staining. Antigen retrieval was performed in Tris-EDTA buffer at pH 9 for four minutes in a pressure cooker set to low pressure. Tissue sections were incubated with anti-PyMT antibody (Novus Biologicals, Cat# NB-100-2749, dilution 1:250) overnight at 4°C and with a biotinylated anti-rat antibody for 1 hour at room temperature. Vectastain Elite ABC (Vector, Cat# PK6100) and ImmPACT DAB Peroxidase (Vector, Cat# SK-4105) were used for signal detection and nuclei were counterstained with methyl green.

### mRNA In Situ Hybridization

Fresh, RNase-free FFPE sections were stained with RNAscope multiplex fluorescent reagent kit V2 according to standard manufacturer protocol. Target retrieval was performed using 8 minute incubation in a pressure cooker set to low pressure. Opal 520 and Opal 570 (PerkinElmer) were used at 1:1500 for target visualization.

### Gene expression by RT-qPCR

Total RNA was reverse-transcribed using random primers (Amersham Bioscienes) and results were normalized to 18S RNA to control for varying cDNA concentration between samples. The primer sequences used are 18s forward 5’-GGATGCGTGCATTTATCAGA-3’ and reverse 5’-GGCGACTACCATCGAAAGTT-3’, Lox forward 5’-CGGGAGACCGTACTGGAAGT-3’ and reverse 5’-CCCAGCCACATAGATCGCAT-3’, Loxl2 forward 5’-CACAGGCACTACCACAGCAT-3’ and reverse 5’-CCAAAGTTGGCACACTCGTA-3’, and Tgfb1 forward 5’-TCATGTCATGGATGGTGCCC-3’ and reverse 5’-GTCACTGGAGTTGTACGGCA-3’.

### Atomic force microscopy data acquisition

Atomic force microscopy (AFM) measurements were performed as previously described(2). Briefly, 20µm OCT-embedded frozen human breast tissue or 30µm mouse mammary gland sections were fast thawed by immersion in PBS at room temperature. Next, these sections were immersed in PBS containing phosphatase inhibitors (Roche, Cat# 04906845001), protease inhibitor (Roche, Cat# 04693124001), and propidium iodide (ACROS, Cat# 440300250) and placed on the stage for AFM measurements. AFM indentations were performed using an MFP3D-BIO inverted optical AFM (Asylum Research) mounted on a Nikon TE2000-U inverted fluorescent microscope. Silicon nitride cantilevers were used with a spring constant of 0.06 N m^-1^ and a borosilicate glass spherical tip with 5 µm diameter (Novascan Tech). The cantilever was calibrated using the thermal oscillation method prior to each experiment. The indentation rate was held constant within each study but varied between 2-20 μms^-1^ with a maximum force of 2 nN between studies. Force maps were obtained as a raster series of indentations utilizing the FMAP function of the IGOR PRO build supplied by Asylum Research. Elastic properties of ECM were reckoned using the Hertz model. A Poisson’s ratio of 0.5 was used in the calculation of the Young’s elastic modulus.

### Western blotting

Snap frozen tissues were ground while frozen and lysed in 2% SDS containing protease and phosphatase inhibitor. Samples were boiled for 5 minutes (95°C) and loaded onto the SDS-polyacrylamide gel, and protein was separated at 110 constant volts. The protein was transferred onto a pre-wet polyvinylidene difluoride (PVDF) membrane (100% methanol, 1 minute) at 250 mA for 2 hours. The PVDF membrane was rinsed with TBST and non-specific binding was blocked with 5% nonfat dry milk dissolved in TBST. The membrane was then incubated with the primary antibody overnight at 4°C, washed with TBST, incubated with horseradish-peroxidase conjugated secondary antibody (1 hour, room temperature; dilution 1:5000), washed with TBST, and detected with the chemiluminescence system Quantum HRP substrate (Advansta #K-12042). Quantification was performed using gel densitometry in ImageJ. Primary antibodies used are anti-LOX (1:1000, Abcam Cat# ab174316) and anti-E-cadherin (1:1000, Cell Signlaing Cat#3195).

### Flow cytometry

Mouse tissue was harvested and chopped with a razor blade. Chopped tissue was digested in 100 U/mL Collagenase Type 1 (Worthington Biochemical Corporation, Cat# LS004196), 500 U/mL Collagenase Type 4 (Worthington Biochemical Corporation, Cat# LS004188), and 200 µg/mL DNase I (Roche, Cat# 10104159001) while shaking at 37°C. Digested tissue was filtered using a 100 µm filter to remove remaining pieces. Red blood cells were lysed in ammonium-chloride-potassium buffer and remaining cells were counted. Cells were stained with fluorophore-conjugated primary antibodies for 30 minutes on ice and subsequently stained with a viability marker. Antibodies used for staining were anti-mouse CD24-PE (BD Pharmingen, Cat# 553262), anti-mouse TER-119-APC (BioLegend, Cat# 116212), anti-mouse CD45-APC (BioLegend, Cat# 103112), anti-mouse CD29-AF700 (BioLegend, Cat# 102218), anti-mouse Ly6G-BV421 (BioLegend, Cat# 127628), anti-mouse F4/80-BV510 (BioLegend, Cat# 123135), anti-mouse CD29-AF488 (BioLegend, Cat# 102212), anti-mouse CD140a-PE (BioLegend, Cat# 135906), anti-mouse CD31-APC (BioLegend, Cat# 102410), anti-mouse CD11c-BV605 (BD Pharmingen, Cat# 563057), anti-mouse CD24-BV650 (BD Pharmingen, Cat# 563545), anti-mouse CD11b-PerCP-Cy5.5 (eBiosciences, Cat# 45-0112-82), anti-mouse CD45-AF700 (BioLegend, Cat# 103128), anti-mouse Ly6C-BV711 (BioLegend, Cat# 128037), anti-mouse MHCII-PE-Cy7 (BioLegend, Cat# 107630), and Zombie NIR Fixable Viability Dye (BioLegend, Cat# 423105). Cells were then analyzed on a flow cytometer.

### Patient gene expression analysis

For the stroma and epithelium specific gene expression analysis, the breast cancer datasets from Finak *et al.* 2008 and Gruosso *et al.* 2019 have been used. Briefly, whole Human Genome 44 K arrays (Agilent Technologies, product G4112A) were used for stroma and epithelial expression profiles. Details of laser capture microdissection, RNA extraction, labeling, hybridization, scanning and quality filters are described in Finak *et al.,* 2006 and 2008. Briefly, the dataset was normalized using loess (within-array) and quantile (between-array) normalization. Probes were ranked by Inter-quartile range (IQR) values, and the most variable probe per gene across expression data were selected for further analysis. Replicate arrays with a concordance above 0.944 were averaged before assessing differential expression.

An association between PLOD2 and distant metastasis-free survival (DMFS) has been determined using an online tool (http://xena.ucsc.edu) to download GEO data (GSE2034, GSE5327, and GSE7390) from 683 patients analyzed on Affymetrix U133A platform as described in Yau et al.(46). Patients have been excluded from analyses if their molecular subtyping of ER/HER2 status and PAM50 did not align: ER+/HER2-must always be luminal, ER+ or-/HER2+ must be HER2+, and ER-/HER2-must always be basal-like. PLOD2 expression levels have been divided based on the median for each tumor subtype: ER+/HER2-(low n=157; high=157), ER- or +/HER2+ (low n=36; high=37), and ER-/HER2-(low n=66; high=67). All statistical analyses were done using GraphPad Prism Version 6.01: Kruskal-Wallis one-way ANOVA test was applied to assess the relationship in PLOD2 expression levels among tumor subtypes and log rank P value (Mantel-Cox) tests for DMFS curves.

An association between PLOD2 gene expression and relapse-free survival (RFS) has been determined using an online tool (http://kmplot.com/analysis/) from 1,809 patients analyzed on Affymetrix platform (HGU133A and HGU133+2 microarrays)(47). Affymetrix ID 202619 or 202620 were used for PLOD2 probes (2014 version) in these analyses. All breast cancer patients in this database were included regardless to lymph node status, TP53 status, or grade. No restrictions were placed in term of patient treatment. PLOD2 expression levels have been divided based on the median for each tumor subtype: ER+/PR+/HER2-(low n=170; high=169), ER-/PR-/HER2+ (low n=58; high=57), and ER-/PR-/HER-(low n=128; high=127). Hazard ratio (and 95% confidence intervals) and log rank P values were calculated and displayed once the data were plotted using the online tool.

The cBioPortal for Cancer Genomics was used to determine the levels of LOX, PLOD2, and LOXL2 gene expression in breast cancer patients segregated by ER and HER2 status(59, 60). ER, PR, and HER2 status were determined by gene expression levels. Samples positive for both ER/PR and HER2 overexpression were excluded from subtype analysis. The cBioPortal was also used to assess gene expression associations of LOX, PLOD2, and LOXL2 with HIF1A, CCL2, CD163, and CD68. All data accessed via cBioPortal are from the 1904 patients in the METABRIC dataset analyzed for gene expression by Illumina human v3 microarray(61). All 1904 samples were included in correlation analyses.

### Statistical analysis

GraphPad Prism Version 6.01 was used to perform all statistical analyses with the exception of LH2 IHC and RFS correlations with PLOD2. Statistical significance was determined using the appropriate tests as noted in the figure legends or method section.

### LH2 IHC and prognostic analyses

#### Study population

The female Malmö Diet and Cancer Study (MDCS) cohort consists of women born 1923– 1950)(62, 63). Information on incident breast cancer is annually retrieved from the Swedish Cancer Registry and the South Swedish Regional Tumor Registry. Follow-up until December 31, 2010, identified a total of 910 women with incident breast cancer, the following conditions excluded patients: 1) with in situ only cancers (n=68), 2) who received neo-adjuvant treatments (n=4), 3) with distant metastasis at diagnosis (n=14), 4) those who died from breast cancer-related causes ≤ 0.3 years from diagnosis (n=2), and finally 5) patients with bilateral cancers (n=17). In addition, one patient who declined treatment for four years before accepting surgery was excluded. Patient characteristics at diagnosis and pathological tumor data were obtained from medical records. Information on cause of death and vital status was retrieved from the Swedish Causes of Death Registry, with last follow-up December 31st, 2014. Ethical permission was obtained from the Ethical Committee at Lund University (Dnr 472/2007). All participants originally signed a written informed consent form.

#### Tumor evaluation

Tumor samples from incident breast cancer cases in MDCS were collected, and a tissue microarray (TMA) including two 1-mm cores from each tumor was constructed (Beecher, WI, USA). Within the study population (N=910), tumor tissue cores were accessible from 718 patients. Four-μm sections dried for one hour in 60^°^C were automatically pretreated using the Autostainer plus, DAKO staining equipment with Dako kit K8010 (Dako, DK). A primary mouse monoclonal Lysyl Hydroxylase 2 (LH2) antibody (Origene; Cat# TA803224, dilution 1:150) was used for the immunohistochemical staining.

TMA cores were analyzed by a cohort of 4 anatomic pathologists (ACN, AC, JG, AN) using the PathXL digital pathology system (http://www.pathxl.com, PathXL Ltd., UK) blinded to all other clinical and pathologic variables. Immunohistochemistry for LH2 was assessed separately for stromal and neoplastic epithelial components of the tumors. Stromal LH2 staining was assessed with the semi-quantitative H-score which combines intensity and proportion positive assessments into a continuous variable from 0-300(64). Cellular stromal components were assessed (including fibroblasts, macrophages, endothelial cells, adipocytes, and other stromal cell types) while areas of significant lymphocytic infiltrate were specifically excluded from the percent positive estimation. Neoplastic epithelial LH2 staining intensity was scored 0-3+ based on the predominant intensity pattern in the tumor — invasive tumor cells did not display significant intra-tumoral heterogeneity of LH2 staining within each core. Verification of inter-observer reproducibility for the H-score was established in a training series of 16 cases evaluated by all study pathologists to harmonize scoring. Inter-observer agreement in the training set was very high, evaluating the IHC scores both as continuous variables (Pearson correlation coefficients ranging from 0.912-0.9566, all p values < 0.0001), and after transformation into categorical data (negative, low, and moderate/high; weighted kappa coefficients ranging from 0.673-0.786). In addition, 50 cases of the study cohort were evaluated blindly by two pathologists to confirm data fidelity; the Pearson correlation coefficient = 0.7507 (p = 5.7 E-05), considered a strong level of agreement.

After exclusion of cases for which LH2 was not evaluable on the TMA, H-scores for 505 patients were included for statistical associations with clinicopathologic features and patient outcome. Each patient was represented by two cores, and TMA core 1 and core 2 were merged into a joint variable favoring the highest stromal LH2 H-score or epithelial LH2 intensity because we predict that higher H-score would drive patient outcome in accordance with our gene expression data demonstrating high LH2 expression correlated with poor outcome. The Pearson correlation coefficient between cores = 0.647, demonstrating moderate agreement among the stromal LH2 H-scores for the two cores. In cases with only one TMA core providing a LH2 score, the expression of this core was used. Further, the joint stromal LH2 variable was categorized into tertiles based on the study population with valid LH2 annotation (N=505). The lowest tertile of LH2 H-scores were defined as scores between 0 and less or equal to 120 (N=171), the intermediate H-score as above 120 and equal or less than 230 (N=188), and the highest stromal LH2 score as above 230 (N=146).

#### Statistical analyses for LH2 IHC analyses

Patient and tumor characteristics at diagnosis in relation to stromal LH2 expression were categorized and presented as percentages. Continuous variables are presented as the mean and min/max. The associations between LH2 expression and grade or tumor size, respectively, were analyzed through linear-by-linear association. The association between LH2 expression and prognosis was examined using breast cancer-specific mortality as endpoint, which was defined as the incidence of breast cancer-related death. Follow-up was calculated from the date of breast cancer diagnosis to the date of breast cancer-related death, date of death from another cause, date of emigration or the end of follow-up as of December 31st, 2014. Main analyses included the overall population; additional analyses were performed in subgroup analyses stratified by estrogen receptor (ER) or axillary lymph node involvement (ALNI) status. The prognostic impact of stromal LH2 expression was analyzed through Cox proportional hazards analyses, which yielded hazard ratios (HR) and 95% confidence intervals (CI) for crude models, and multivariate models adjusted for age at diagnosis (model 1) and tumor characteristics ER (dichotomized, cut-off 10% stained nuclei), ALNI (none or any positive lymph node involvement), histological grade (Nottingham grade I-III), and tumor size (dichotomized using cut-off 20 mm). Kaplan-Meier curves including the LogRank test indicated LH2 status to particularly impact the first 10 years after diagnosis and survival variables constructed to capture these effects were used in Cox regression models investigating the effects during the first post-diagnostic decade. All statistical analyses were performed in SPSS version 22.0 (IBM).

## Author Contributions

V.M.W., K.C.H., O.M., A.S.B., and A.P.D. conceived the project, prepared figures and wrote the manuscript. A.S.B, T.P., T.N., and K.C.H. developed the xAAA method and A.S.B performed all LC-MS and LC-PRM experiments. J.N.L generated the TetO_mLOX mouse model. O.M. designed and conducted *in vivo* experiments using inducible LOX overexpression models. B.R. and L.C. designed and conducted CSF1 blocking antibody mouse experiment. O.M. and A.P.D. performed and quantified immunofluorescence, H&E, PS and SHG imaging and analyses on mouse tissue samples. I.A., A.P.D., and J.M.B performed AFM on human or mouse tissue specimens. I.A. performed SHG imaging on human tissue. E.S.H, and P.K. provided human breast tumor biopsies for xAAA. B.R. and L.C. designed and conducted immunoprofiling on human breast tumor via flow cytometry. O.M. and A.P.D. performed all gene expression analyses with the exception of Fig. 5g-j. T.G, H.K. and M.P. performed gene expression analyses gene expression in microdissected epithelial and stromal compartments of human invasive breast carcinomas. O.M. and A.P.D. designed and conducted *in vivo* experiments using minoxidil treatment. S.B. established and managed MDCS cohort used for LH2 IHC. Z.W. and S.B. performed LH2 IHC. A.C.N. designed scoring schemes for stromal and neoplastic epithelial LH2 IHC, and A.C.N, A.N., J.G., and A.C. scored all human biopsies. S.B., O.B., A.C.N., O.M., and V.M.W analyzed and interpreted clinical data from LH2 scores.

## Acknowledgements

We thank J. Northcott for writing the ImageJ Macro, L. Korets for mouse husbandry and N. Korets for histology support, as well as K. Lövgren and S. Baker for LH2 immunostaining on patient biopsies. The work was supported by investigator grants through the US National Cancer Institute R33 CA183685 (K.C.H & V.M.W) and R01CA192914 and CA174929 to (V.M.W), and R01CA222508-01 to (V.M.W. and E.S.H.), as well as US DOD Breast Cancer Research Program (BCRP) grant BC122990 (V.M.W). Trainee support was provided by US DOD BCRP grant BC130501 (O.M.), US NIH grants TL1 TR001081 & US NIH T32 HL007171 (A.S.B), and US NIH T32 grant CA 108462 (O.M.). Funding from Eastern Star Scholar-Minnesota Masonic Cancer Center (A.C.N.), the Swedish Research Council (S.B. & C.H.) and US NIH R01 CA057621 (Z.W.) also supported the work.

