## Supplementary Material for "Inflammation promotes tumor aggression by stimulating stromal cell-dependent collagen crosslinking and stromal stiffening"

**Suppl Table 1**

| xAA Standard | Letter Code | Elemental Composition | RT (min) | Mass (Da) | Accuracy (ppm) | LLOD (fmole) | $R^2$ | LLOQ (fmole) | Structure |
|---|---|---|---|---|---|---|---|---|---|
| Lysinonorleucine | LNL | C12H26N3O4+ | 5.69 | 276.19 | 1.5 | 75 | 0.96 | 225 | 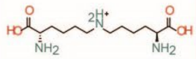 |
| Dihydroxy lysinonorleucine | DHLNL | C12H26N3O6+ | 7.16 | 308.18 | 2 | 125 | 0.95 | 375 | 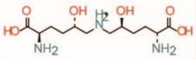 |
| deoxy Lysyl pyridinoline | dPyr | C18H29N4O7+ | 8.14 | 413.2 | 4.5 | 250 | 0.98 | 750 | 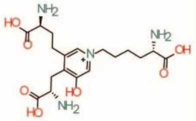 |
| Desmosine* | Des | C24H40N5O8+ | 10.14 | 526.29 | 0.9 | 156 | 0.92 | 468 | 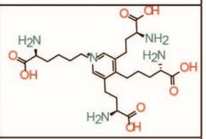 |
| Isodesmosine* | iDes | C24H40N5O8+ | 10.14 | 526.29 | 0.9 | 156 | 0.92 | 468 | 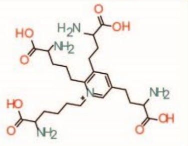 |

**Supplementary Table 1: Summary of crosslinked amino acid standard characterization by mass spectrometry.** Serial dilutions of crosslinked amino acid standards were prepared in the background of *E. coli* hydrolysates and the lowest limit of detection (LLOD) and the lowest limit of quantification (LLOQ) were determined on a QExactive mass spectrometer. The LLOD is defined here as the concentration that is required to produce a signal that is three times the noise level. The LLOQ is the analyte concentration that is required to produce a signal that is three times that of the LLOD. *Denotes isomers that were not resolved.

### Suppl Table 2

| | Distribution of patient and tumor characteristics | | | |
|---|---|---|---|---|
| All n=910 | | | | |
| Tumor in tissue microarray, n (%) | | Yes 718/910 (78.9) | | |
| **Lysyl hydroxylase 2 (LH2) epithelial** expression assessable, n (%) | | Yes, 468/718 (65.2%) | | |
| LH2 neg/weak/moderate/strong | | **LH2 negative** | **LH2 weak** | **LH2 moderate/strong** |
| | | 279 (59.6%) | 112 (23.9%) | 77 (16.5%) |
| Factor | n (%) or mean (min-max) | n (%) or mean (min-max) | n (%) or mean (min-max) | n (%) or mean (min-max) |
| **Age at diagnosis** | 65.5 (45.7-87.3) | 63.9 (48.4-84.7) | 61.7 (46.4-85.6) | 63.9 (45.7-87.3) |
| years (n= 910) | | | | |
| **BMI at baseline (n=910)** | | | | |
| <25 | 466 (51) | 154 (56.8) | 64 (57.1) | 36 (46.8) |
| ≥25 and <30 | 310 (34) | 77 (28.4) | 36 (32.1) | 27 (35.1) |
| >30 | 134 (15) | 40 (14.8) | 12 (10.7) | 14 (18.2) |
| **Tumor size (n= 887)** | | | | |
| ≤20 mm | 637 (72) | 190 (70.6) | 77 (68.8) | 43 (55.8) |
| >20 mm | 250 (28) | 79 (29.4) | 35 (31.3) | 34 (44.2) |
| **ALNI (n=859)** | | | | |
| Negative | 588 (68.5) | 175 (67.6) | 67 (61.5) | 50 (65.8) |
| Positive (≥1 metastatic node) | 271 (31.5) | 84 (32.4) | 42 (38.5) | 26 (34.2) |
| **Grade, NHG (n=860)** | | | | |
| I | 233 (27.1) | 84 (31.7) | 21 (18.9) | 12 (15.8) |
| II | 405 (47.1) | 138 (52.1) | 52 (46.8) | 14 (18.4) |
| III | 222 (25.8) | 43 (16.2) | 38 (34.2) | 50 (65.8) |
| **ER status (n=784)** | | | | |
| Positive (>10%) | 690 (88.0) | 230 (93.5) | 93 (87.7) | 44 (63.8) |
| Negative (<10%) | 94 (12.0) | 16 (6.5) | 13 (12.3) | 25 (36.2) |
| **HER2 status (n=609)** | | | | |
| Negative | 556 (91.3) | 161 (89.9) | 71 (86.6) | 50 (89.3) |
| Positive | 53 (8.7) | 18 (10.1) | 11 (13.4) | 6 (10.7) |
| **Ki67 (n=655)** | | | | |
| Low (≤10%) | 434 (66.3) | 143 (71.5) | 58 (61.7) | 18 (32.7) |
| High (>10%) | 221(33.7) | 57 (28.5) | 36 (38.3) | 37 (67.3) |
| **Molecular subtypes (n=639)** | | | | |
| ER+/HER2- | 536 (83.9) | 169 (88.0) | 76 (80.9) | 29 (53.7) |
| HER2+ | 53 (8.3) | 5 (4.4) | 11 (11.7) | 6 (11.1) |
| TNBC | 50 (7.8) | 3 (2.7) | 7 (7.4) | 19 (35.2) |

**Supplementary Table 2: Characterization of breast cancer patients used to develop neoplastic epithelial LH2 H-score.**

Suppl Table 3

| | Distribution of patient and tumor characteristics | | | |
|---|---|---|---|---|
| All n=910 | | | | |
| Tumor in tissue microarray, n (%) | | | Yes 718/910 (78.9) | |
| **Lysyl hydroxylase 2 (LH2) stromal** expression assessable, n (%) | | | Yes, 505/718 (70.3%) | |
| LH2 low/intermediate/high | | **LH2 low** | **LH2 intermediate** | **LH2 high** |
| | | 171 (33.9%) | 188 (37.2%) | 146 (28.9%) |
| Factor | n (%) or mean (min-max) | n (%) or mean (min-max) | n (%) or mean (min-max) | n (%) or mean (min-max) |
| **Age at baseline** | 56.4 (44.7-73.0) | 53.7 (44.9-73.0) | 54.2 (44.7-72.7) | 53.6 (45.0-72.8) |
| years (n= 910) | | | | |
| **Age at diagnosis** | 65.5 (45.7-87.3) | 62.6 (48.5-84.7) | 63.4 (45.7-87.3) | 63.7 (46.4-85.6) |
| years (n= 910) | | | | |
| **BMI at baseline (n=910)** | | | | |
| <25 | 466 (51) | 98 (57.3) | 110 (58.5) | 73 (50.0) |
| ≥25 and <30 | 310 (34) | 52 (30.4) | 55 (29.3) | 47 (32.2) |
| >30 | 134 (15) | 20 (11.7) | 23 (12.2) | 26 (17.8) |
| **Tumor size (n= 887)** | | | | |
| ≤20 mm | 637 (72) | 118 (69.8) | 131 (70.1) | 95 (65.1) |
| >20 mm | 250 (28) | 51 (30.2) | 56 (29.9) | 51 (34.9) |
| **ALNI (n=859)** | | | | |
| Negative | 588 (68.5) | 116 (72.5) | 116 (64.8) | 90 (62.5) |
| Positive (≥1 metastatic node) | 271 (31.5) | 44 (27.5) | 63 (35.2) | 54 (37.5) |
| **Grade, NHG (n=860)** | | | | |
| I | 233 (27.1) | 53 (32.3) | 60 (32.3) | 21 (14.7) |
| II | 405 (47.1) | 86 (52.4) | 79 (42.5) | 54 (37.8) |
| III | 222 (25.8) | 25 (15.2) | 47 (25.3) | 68 (47.6) |
| **ER status (n=784)** | | | | |
| Positive (>10%) | 690 (88.0) | 133 (94.3) | 155 (89.1) | 106 (77.9) |
| Negative (<10%) | 94 (12.0) | 8 (5.7) | 19 (10.9) | 30 (22.1) |
| **HER2 status (n=609)** | | | | |
| Negative | 556 (91.3) | 104 (95.4) | 120 (88.9) | 85 (84.2) |
| Positive | 53 (8.7) | 5 (4.6) | 15 (11.1) | 16 (15.8) |
| **Ki67 (n=655)** | | | | |
| Low (≤10%) | 434 (66.3) | 83 (72.2) | 98 (66.7) | 54 (48.2) |
| High (>10%) | 221(33.7) | 32 (27.8) | 49 (26.1) | 58 (51.8) |
| **Molecular subtypes (n=639)** | | | | |
| ER+/HER2- | 536 (83.9) | 105 (92.9) | 118 (81.9) | 73 (68.2) |
| HER2+ | 53 (8.3) | 5 (4.4) | 15 (10.4) | 16 (15.0) |
| TNBC | 50 (7.8) | 3 (2.7) | 11 (7.6) | 18 (16.8) |

**Supplementary Table 3: Characterization of breast cancer patients used to develop stromal LH2 H-score.**

**Suppl Fig 1**

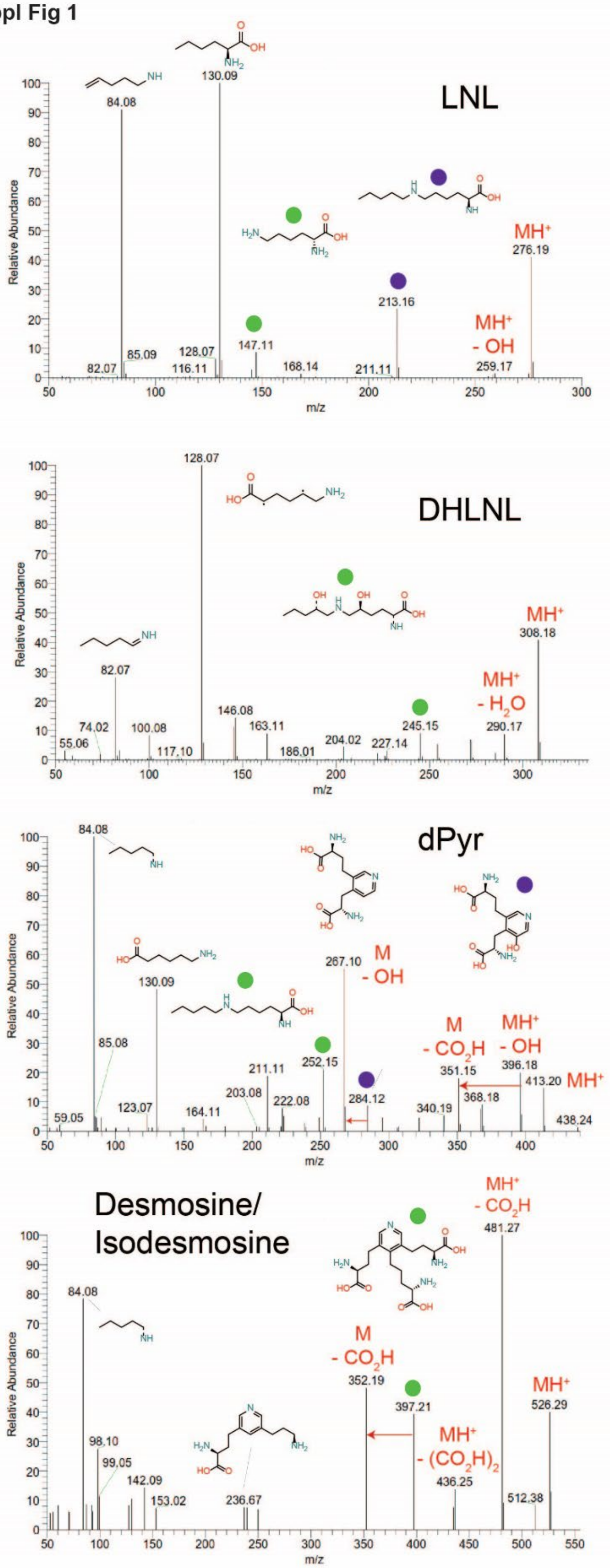

LNL
84.08
130.09
MH+
276.19
213.16
147.11
MH+
- OH
85.09
128.07
82.07
116.11
168.14
211.11
259.17
DHLNL
128.07
MH+
308.18
82.07
MH+
- H2O
146.08
163.11
245.15
290.17
74.02
100.08
55.06
117.10
186.01
204.02
227.14
dPyr
84.08
267.10
M
- OH
130.09
85.08
M
- CO2H
MH+
- OH
211.11
252.15
351.15
396.18
413.20
MH+
203.08
222.08
284.12
368.18
123.07
164.11
340.19
59.05
438.24
Desmosine/
Isodesmosine
MH+
- CO2H
481.27
84.08
M
- CO2H
352.19
397.21
MH+
526.29
MH+
- (CO2H)2
98.10
99.05
142.09
153.02
236.67
436.25
512.38
Relative Abundance
m/z

**Supplementary Figure 1: Partial assignment of MS$^2$ fragmentation spectra of crosslinked amino acid standards.** MS$^2$ fragmentation spectra of commercially available crosslinked amino acid standards with partial assignment of MS$^2$ fragmentation spectra (LNL, **top**; DHLNL, **middle**; dPyr, **bottom**). Protonated forms of precursor ions are denoted by $MH^+$ labels. Colored circles above fragment ions are matched to their suggested fragment ion structure above the full spectra or are listed above the ion if space permits.

Suppl Fig 2

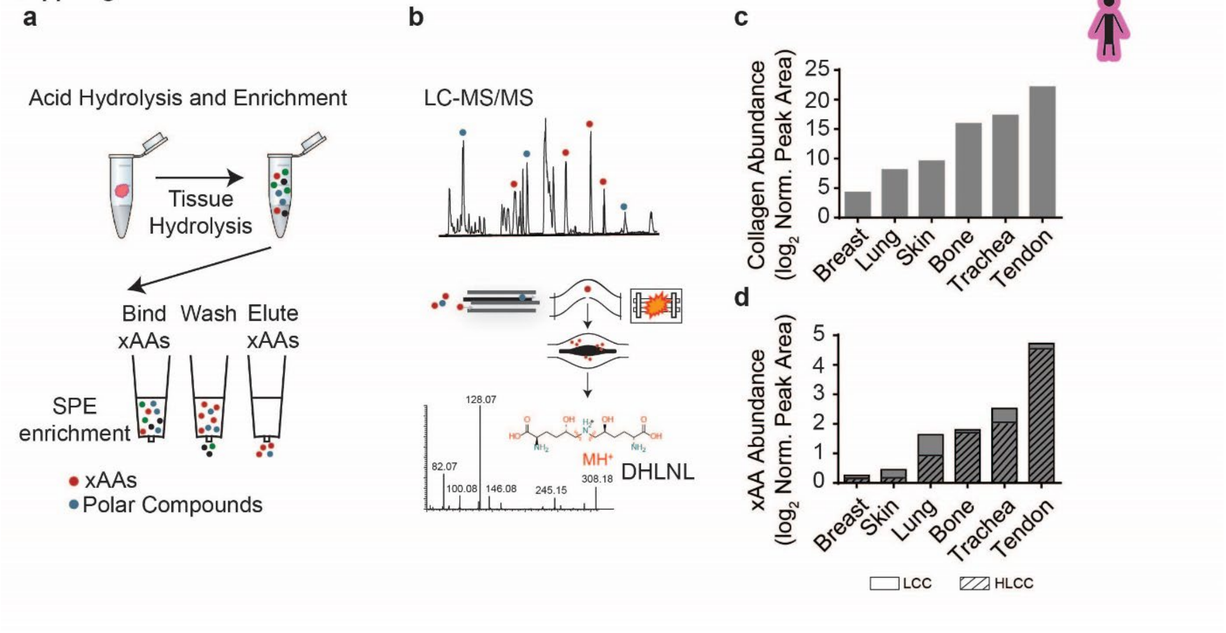

**Supplementary Figure 2: Optimized crosslinking amino acid analysis (xAAA) enables accurate measurement of collagen state across a diverse range of tissue types** (**a**) xAAA workflow schematic. Clinical specimens are hydrolyzed and enriched by solid phase extraction (SPE). (**b**) The enriched hydrolysate is analyzed by LC-SRM on a hybrid quadrupole orbitrap instrument. $MS^2$ spectra is used to accurately identify xAAs such as dihydroxy lysinonorleucine (DHLNL) (**c**) Bar graphs showing quantification of tissue collagen and (**d**) total crosslinked amino acids (xAA) measured in human breast, lung, skin, bone, trachea and tendon (pooled *n* = 3 each tissue). The calculated amino acid crosslink values are normalized to total tissue collagen content (hydroxyproline abundance) which is calculated based on wet tissue weight. The final values have been plotted as relative abundance based on peak area. Bar graphs depict relative abundance of Lysine derived-collagen crosslink (LCC) and hydroxylysine-derived collagen crosslink (HLCC) species.

**Suppl Fig 3**

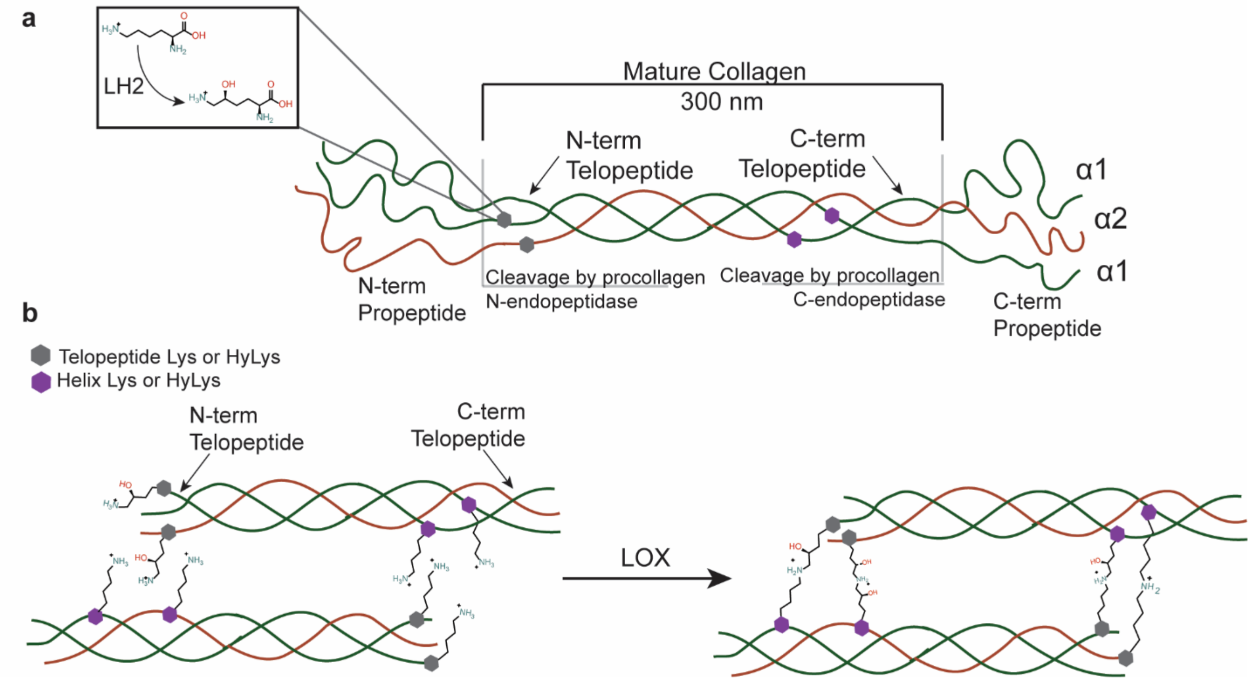

**Supplementary Figure 3: Schematic diagram of lysine hydroxylation and crosslinking in fibrillar collagen.** (**a**) Schematic of a mature fibrillar collagen fiber. N- and C- terminal telopeptides are hydroxylated by lysyl hydroxylase 2 (LH2). N- and C- terminal propeptides are cleaved by procollagen endopeptidases to form the mature collagen fiber (300 nm). (**b**) Lysine (Lys) and the hydroxylysine (Hyl) residues in the telopeptide region of mature collagen are targeted by the crosslinking enzyme lysyl oxidase (LOX), which forms reactive aldehyde groups that spontaneously undergo aldol condensation reactions to form covalent collagen crosslinks.

**Suppl Fig 4 Lysine aldehyde ($Lys^{ald}$) and hydroxylysine aldehyde ($Hyl^{ald}$) collagen crosslinking pathways [7, 17]**

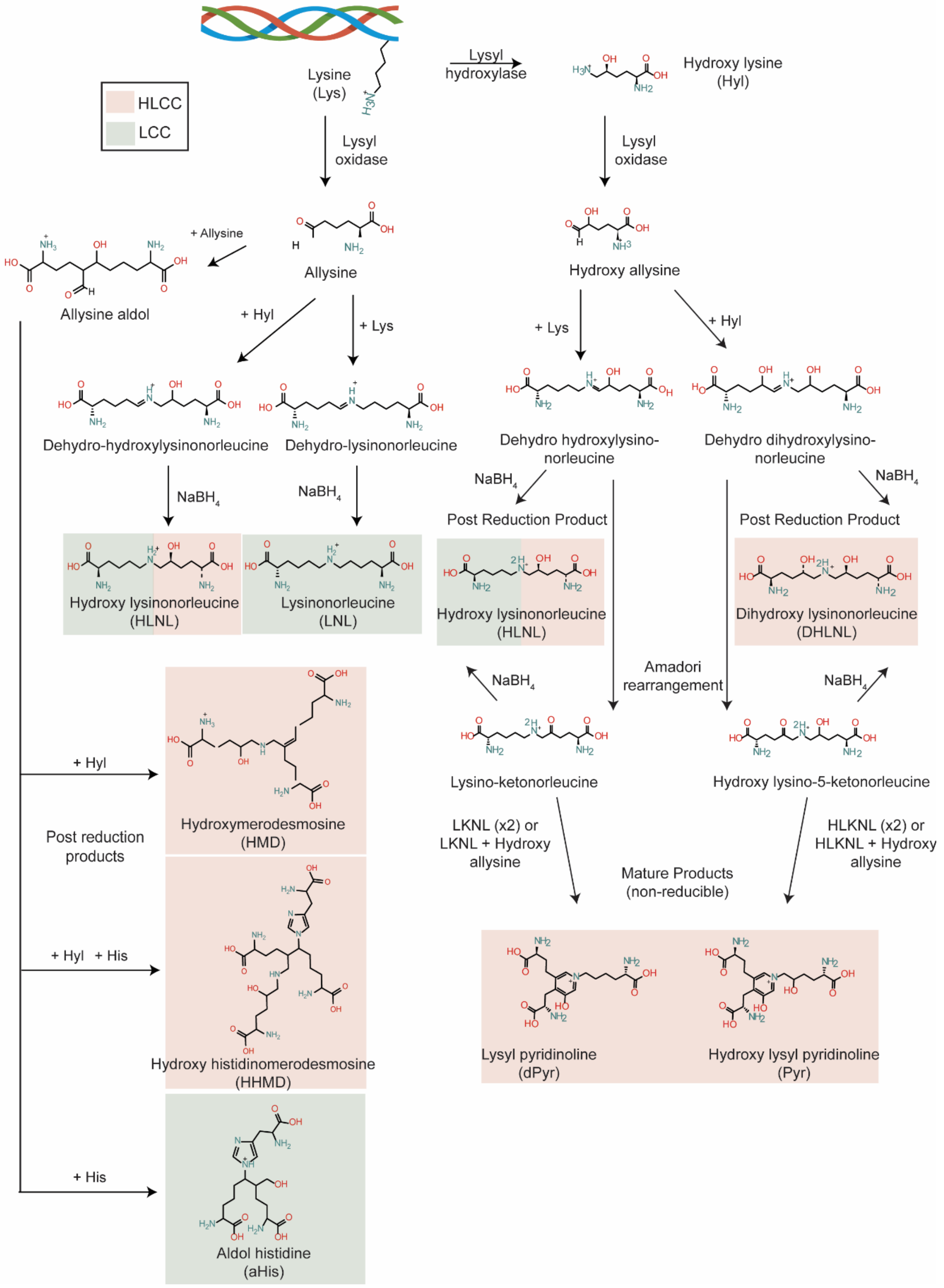

**Supplementary Figure 4: Lysine aldehyde ($Lys^{ald}$) and hydroxylysine aldehyde ($Hyl^{ald}$) crosslinking pathway.** Lysyl oxidase modifies Lys residues to form allysine ($Lys^{ald}$), which spontaneously reacts with Lys and Hyl residues in the helical region to form the Schiff bases dehydro-hydroxylysinonorleucine (deH-HLNL) dehydro-lysinonorleucine (deh-HLNL). These crosslinks can be reduced with $NaBH_4$ to form LNL and HLNL. The mature products of these crosslinks are currently unknown. Allysine can combine with an additional allysine residue to form allysine aldol. Allysine aldol can form the trivalent crosslink hydroxyl merodesmosine, aldol histidine, or the tetravalent crosslink histidino-hydroxymerodesmosine (HHMD) (only post-reduction products shown) through aldol condensation reactions with Hyl or histidine (His), or a combination of the two. Red and green shading denotes lysine-derived collagen crosslinks (LCC) or hydroxyl lysine-derived collagen crosslinks (HLCC)[22]. Telopeptide lysine residues are modified by lysyl hydroxylase. Lysyl oxidase modifies Hyl residues to hydroxyl allysine ($Hyl^{ald}$), which spontaneously reacts with Lys and Hyl residues to form the Schiff bases dehydro-dihydroxylysinonorleucine (deH-DHLNL) and dehydro-hydroxylysinonorleucine (deh-HLNL). They then undergo Amadori rearrangements to form hydroxylysino-keto-norleucine (HLKNL) or lysine-keto-norleucine (LKNL), respectively. These crosslinks can be reduced with $NaBH_4$ to form LNL and DHLNL. Mature crosslink products (Pyr and dPyr) are formed from the reaction of lysine ketonorleucine (LKNL) or hydroxyl lysinoketonorleucine (HKLNL) with hydroxyl allysine. Red and green shading denotes lysine-derived collagen crosslinks (LCC) or hydroxyl lysine-derived collagen crosslinks (HLCC)[22].

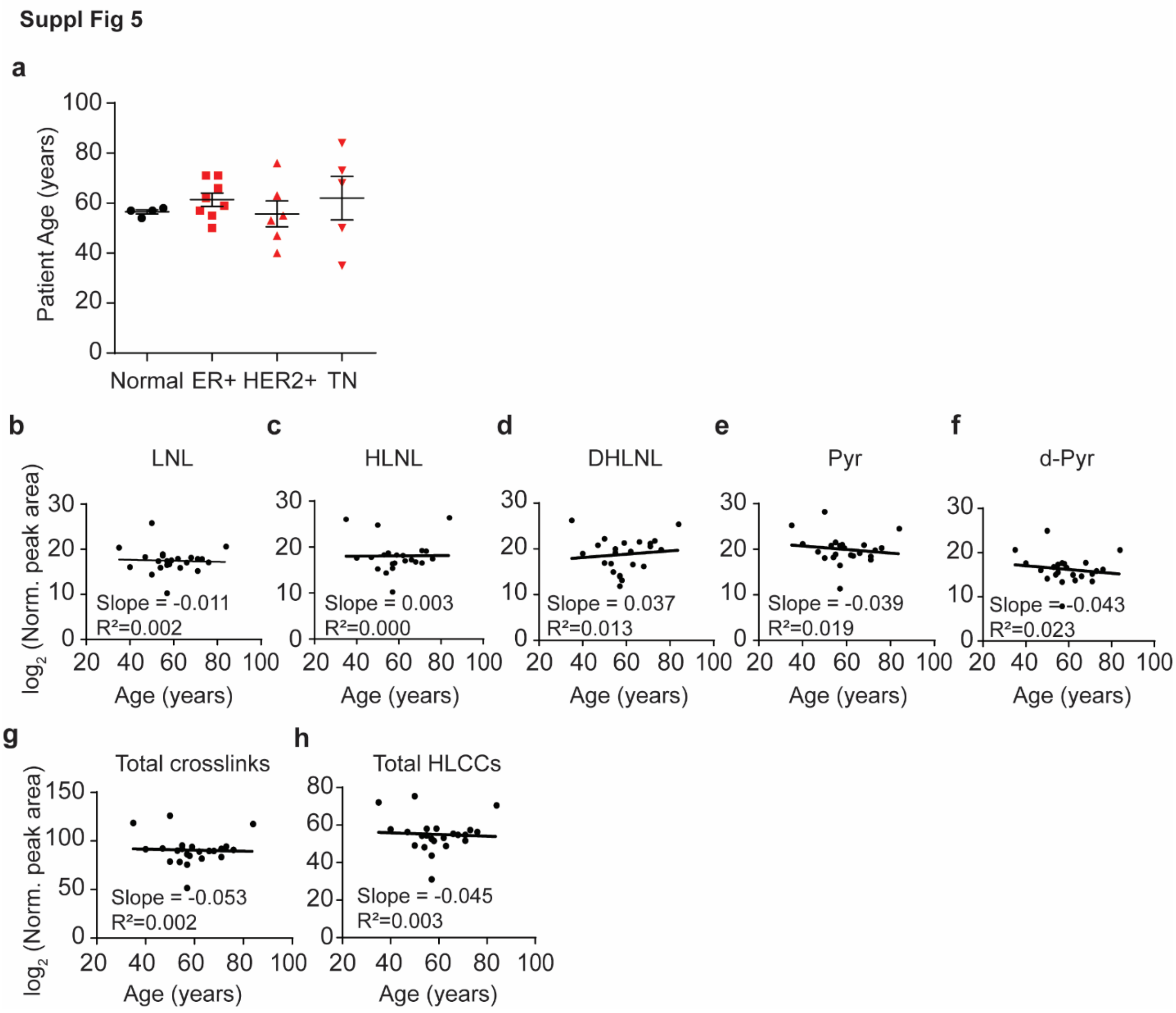

**Supplementary Figure 5: Abundance of LOX-mediated crosslinks does not correlate with patient age.** (**a**) Scatter plot showing the mean and SEM of patient age for normal and tumor tissue samples by subtype. (**b-f**) Scatter plots showing the levels of each LCC and HLCC crosslink measured in normal and tumor breast tissues versus patient age. The total abundance of crosslinks (**g**) was calculated by summing all individual crosslinks and the total tissue HLCC abundance (**h**) was calculated by summing DHLNL, Pyr, and d-Pyr. All crosslink values are normalized to total collagen content (i.e. hydroxyproline abundance) and wet tissue weight and are plotted as $\log_2$ transformed normalized peak areas from LC-MS data. The best fit line and its slope and $r^2$ value are displayed on each plot.

Suppl Fig 6

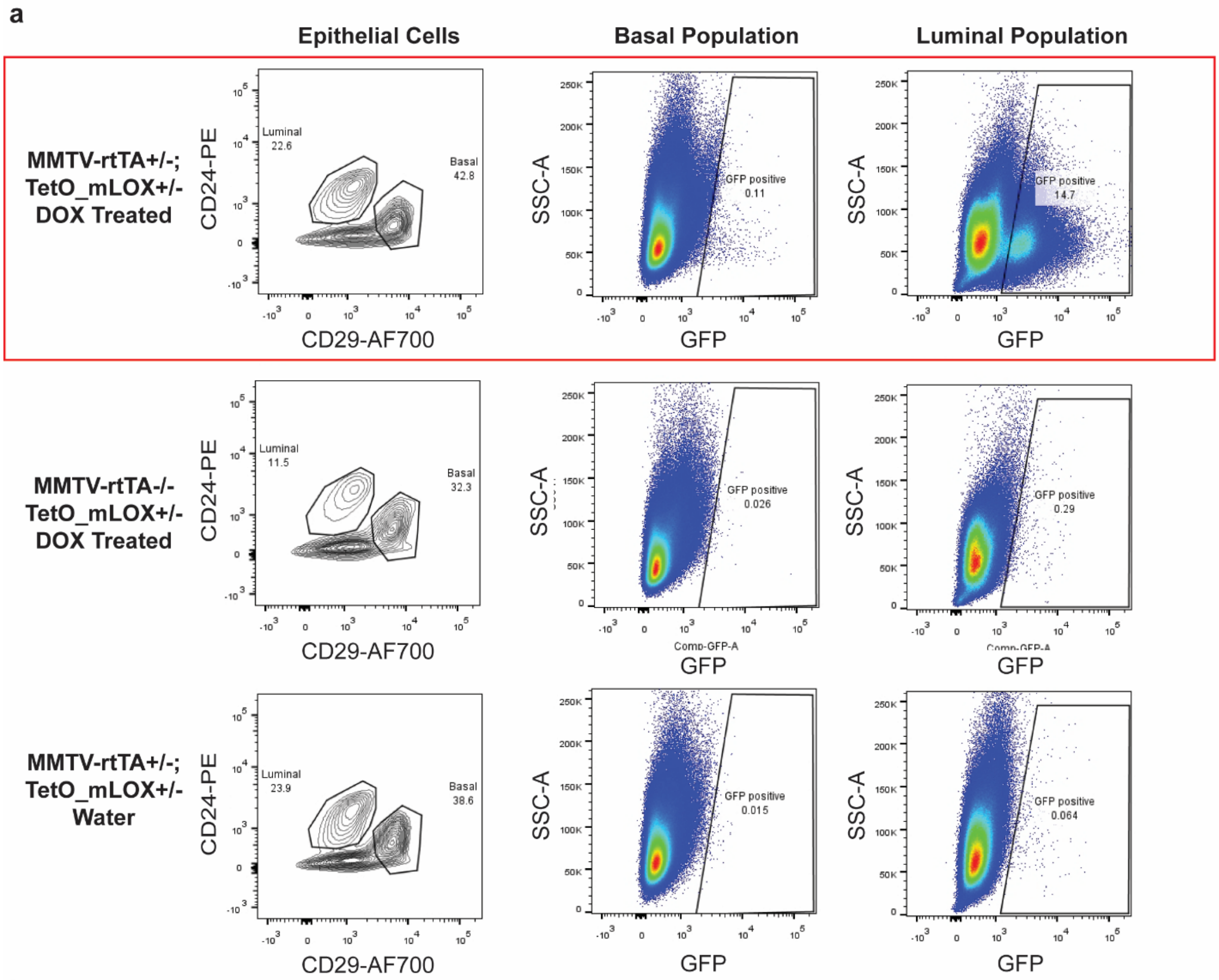

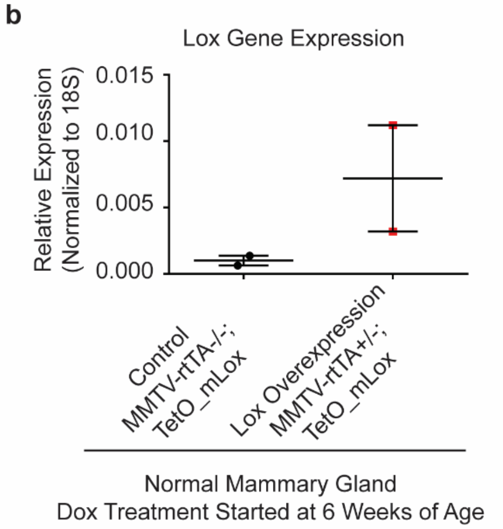

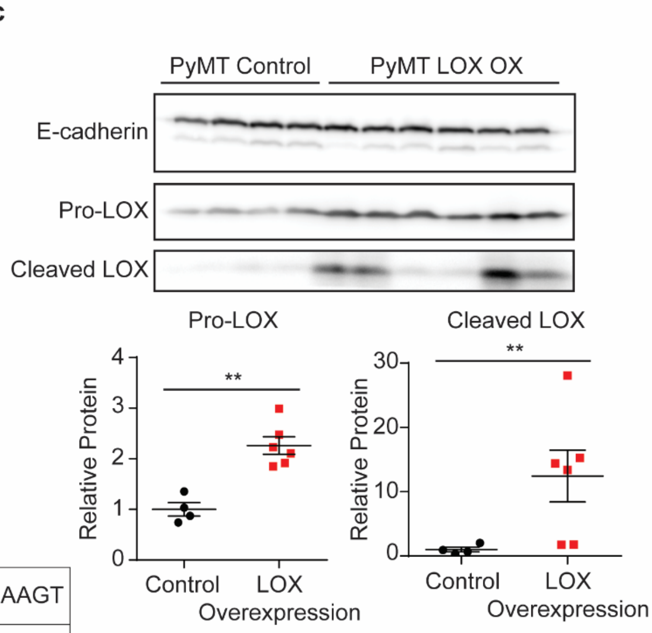

Lox Primers for qPCR:

| | | |
|---|---|---|
| Mouse | Rev | CGGGAGACCGTACTGGAAGT |
| Mouse | For | CCCAGCCACATAGATCGCAT |

**Supplementary Figure 6: Validation of LOX overexpression in doxycycline (DOX) treated MMTV-rtTA$^{+/-}$; TetO_mLox$^{+/-}$ mice.** (**a**) Representative dot plots (n = 2) of GFP expression in mouse mammary epithelial cells analyzed by flow cytometry. GFP and mLOX are encoded on the same mRNA transcript with independent translation initiation sites. (**Top panel**) GFP expression is induced in DOX treated MMTV-rtTA$^{+/-}$; TetO_mLox$^{+/-}$ mice. (**Middle and Bottom panels**) GFP expression is not induced in control mice lacking the MMTV-rtTA promoter (**Middle**) or control mice not treated with DOX (**Bottom**). To select for mammary epithelial cells were gated out cells positive for anti-mouse CD45-APC, anti-mouse CD31-APC, anti-mouse Ter119-APC antibodies. (**b**) Quantification of Lox gene expression in whole mouse mammary gland by RT-qPCR in control MMTV-rtTA$^{-/-}$; TetO_mLox$^{+/-}$ mice (n = 2) and Lox overexpressing MMTV-rtTA$^{+/-}$; TetO_mLox$^{+/-}$ (n = 2) mice. (**c**) Western blot of whole tumor lysate from control and epithelial LOX overexpressing PyMT tumors. Scatter plots with mean ± SEM quantify optical density of each band normalized to E-cadherin. Statistical analyses were performed using Mann-Whitney U test (**$p < 0.01$).

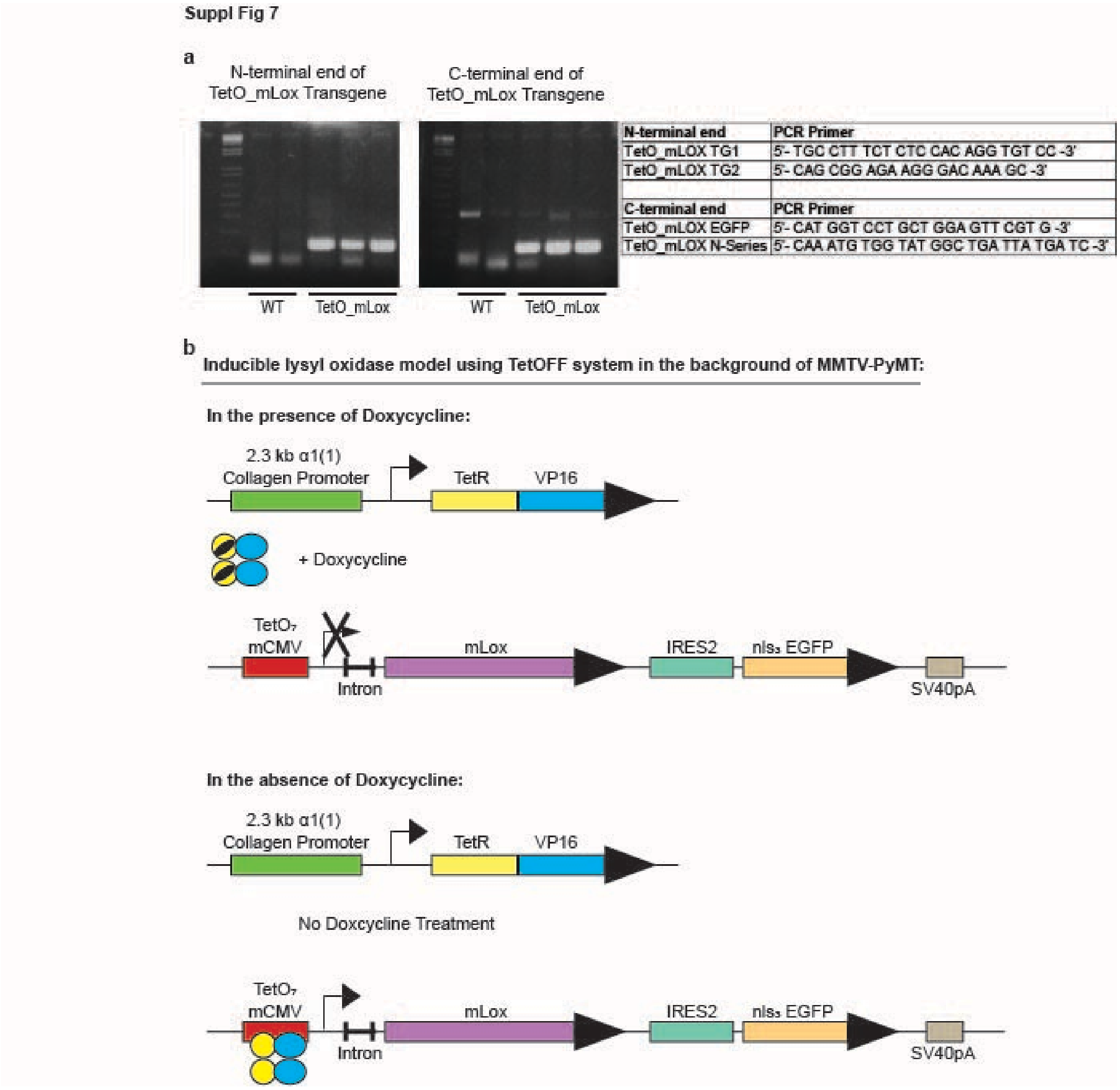

| N-terminal end | PCR Primer |
|---|---|
| TetO_mLOX TG1 | 5'- TGC CTT TCT CTC CAC AGG TGT CC -3' |
| TetO_mLOX TG2 | 5'- CAG CGG AGA AGG GAC AAA GC -3' |
| **C-terminal end** | **PCR Primer** |
| TetO_mLOX EGFP | 5'- CAT GGT CCT GCT GGA GTT CGT G -3' |
| TetO_mLOX N-Series | 5'- CAA ATG TGG TAT GGC TGA TTA TGA TC -3' |

**Supplementary Figure 7: Design of inducible lysyl oxidase expression *in vivo* using a TetOFF system in the background of MMTV-PyMT model.** (**a**) PCR results and sequence of PCR primers of N- and C-terminal ends to confirm TetO_mLox transgene incorporation. (**b**) Diagram of inducible TetOFF system. TetO_mLox transgene is reversibly turned off or on in the presence or absence of doxycycline, respectively.

**Suppl Fig 8**

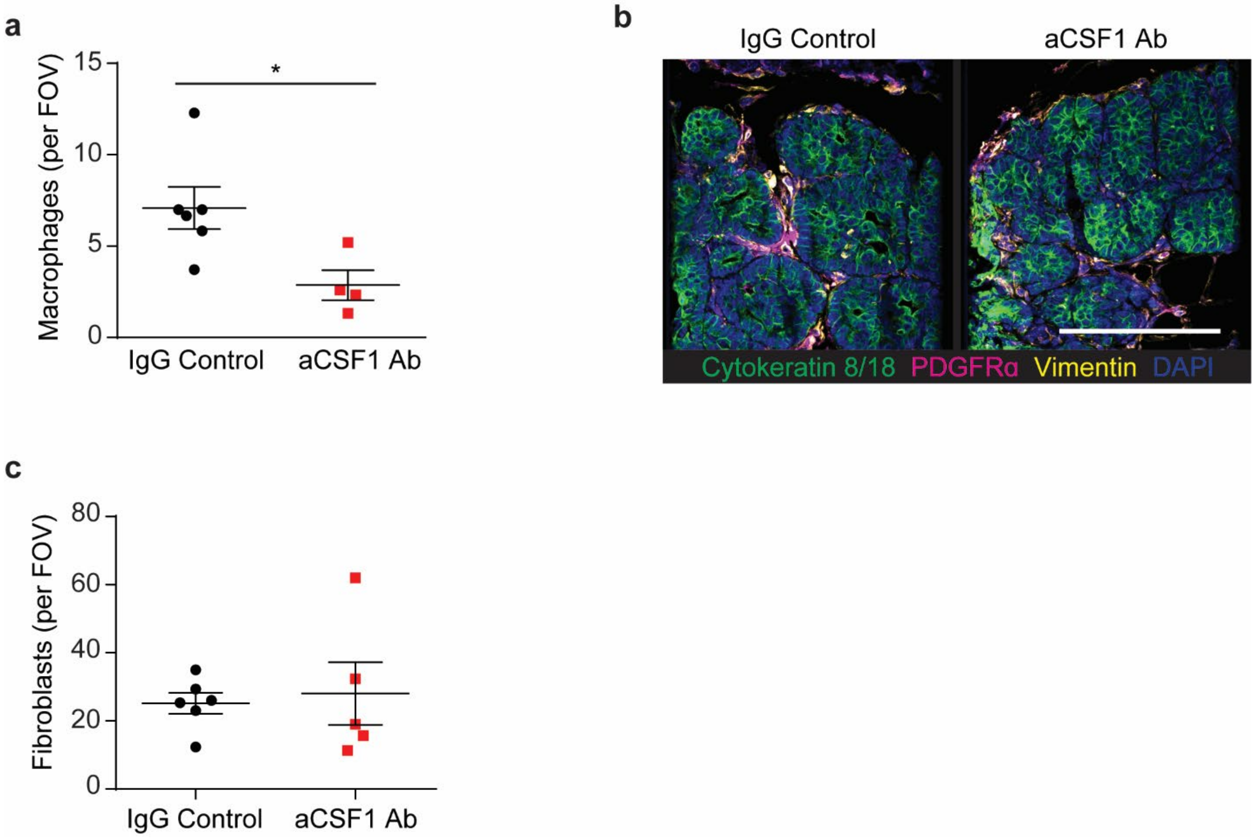

**Supplementary Figure 8: Treatment with anti-CSF1 antibody reduces macrophage accumulation in tumors but has no affect on the fibroblast population.** (**a**) Scatter plot showing mean ± SEM of the number of F4/80 positive cells per field of view in IgG1 treated (n = 6) or anti-CSF1 treated (n = 4) PyMT tumors from 8 week old mice. Statistical analysis was performed using a two-tailed unpaired t-test (*$p < 0.05$). (**b**) Representative images from 8 week old IgG1 treated and anti-CSF1 treated PyMT mice stained for cytokeratin 8/18 (green), PDGFRα (magenta), vimentin (yellow), and DAPI (blue). Scale bar is 100 μm. (**c**) Scatter plot showing mean ± SEM of fibroblasts (vimentin$^+$ and PDGFRα$^+$) per field of view in 8 week old IgG1 treated (n = 6) and anti-CSF1 (n = 5) treated PyMT mice.

Suppl Fig 9

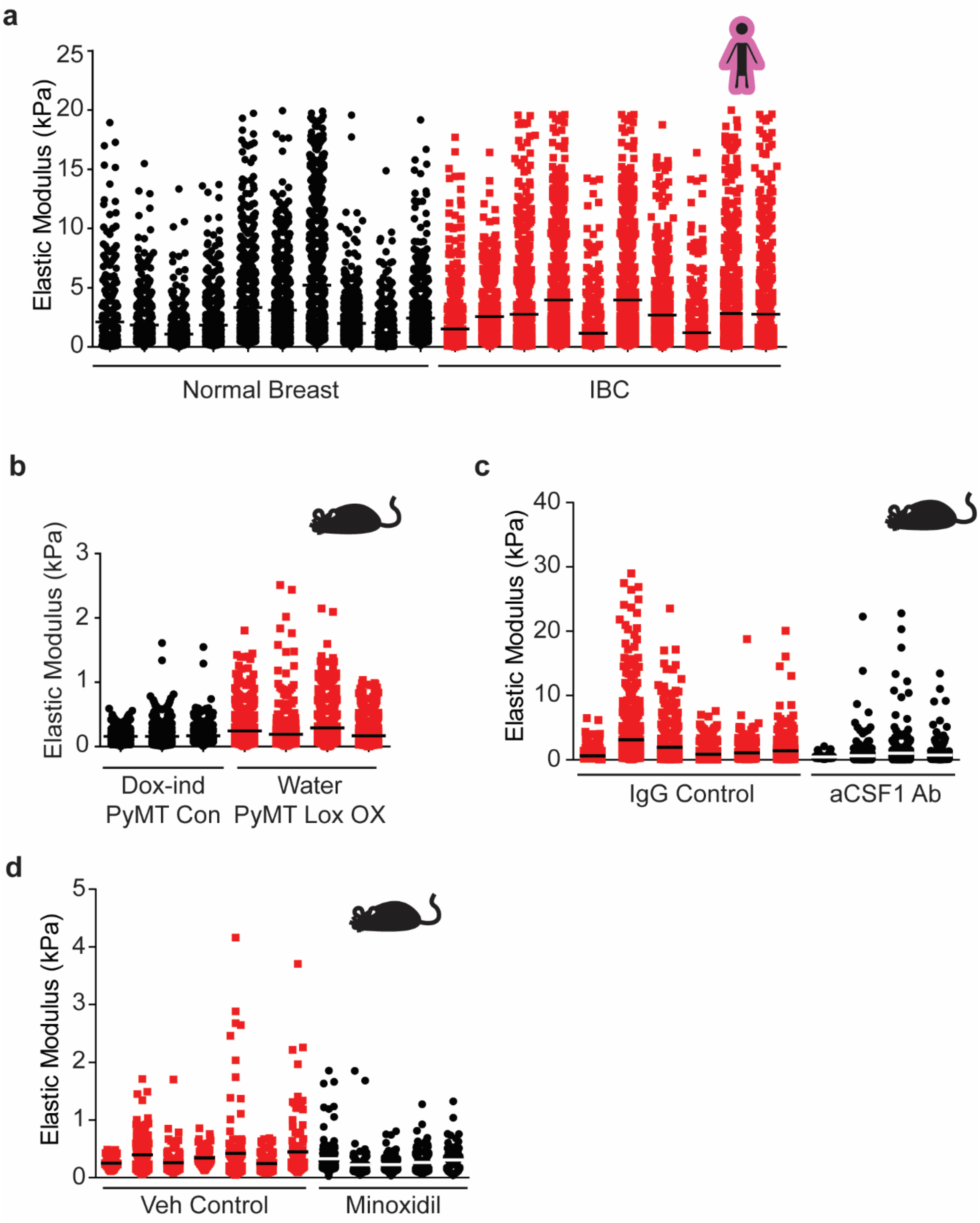

**Supplementary Figure 9: ECM Stiffness for individual specimens.** (**a**) Scatter plot showing mean and individual values of atomic force microscopy (AFM) microindentation measurements of individual human samples categorized as normal breast tissue (black circles) or invasive ductal carcinoma (IDC; red squares). (**b**) Scatter plot showing mean and individual values of

AFM microindentation measurements of individual mouse mammary tumors from doxycycline treated MMTV-PyMT$^{+/-}$; Col1a1-tTA$^{+/-}$; TetO-mLox control mice (DOX-ind PyMT Con; black circles) or water treated mice overexpressing stromal Lox (Water PyMT Lox OX; red squares). (**c**) Scatter plot showing mean and individual values of AFM microindentation measurements of individual mouse mammary gland tumors from 8 week old IgG1 treated (IgG Control; red squares) and anti-CSF1 treated (aCSF1 Ab; black circles) PyMT tumors. (**d**) Scatter plot showing mean and individual values of AFM microindentation measurements of individual mouse mammary gland tumors from PBS vehicle treated (Veh Control; red squares) and minoxidil treated (black circles) PyMT tumors.

Suppl Fig 10

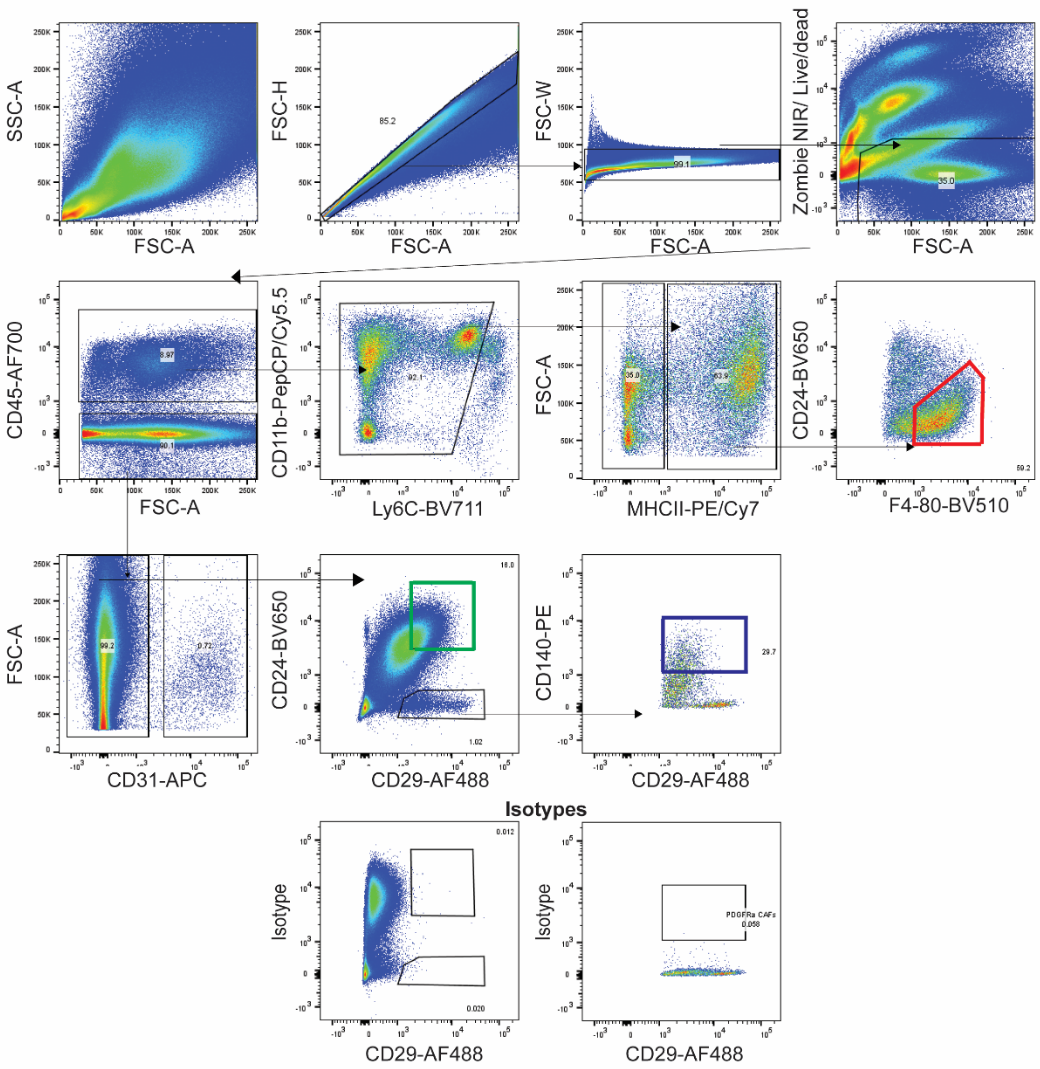

**Supplementary Figure 10: Gating strategy for sorting tumor cells, cancer-associated fibroblasts, and tumor-associated macrophages from PyMT mice via flow cytometry.**

**Suppl Fig 11**

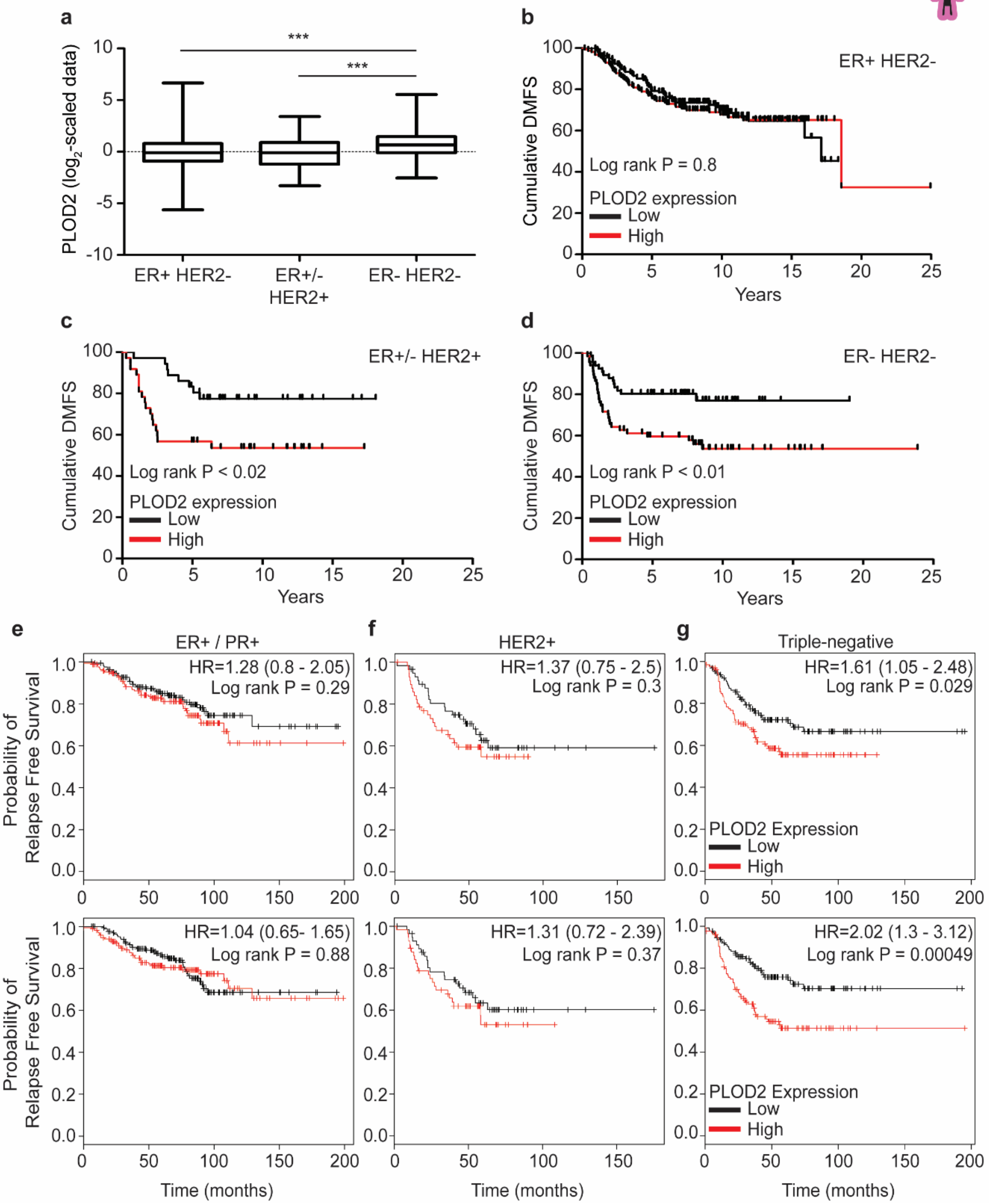
a
PLOD2 (log2-scaled data)
***
***
ER+ HER2-
ER+/- HER2+
ER- HER2-
b
ER+ HER2-
Cumulative DMFS
Log rank P = 0.8
PLOD2 expression
Low
High
Years
c
ER+/- HER2+
Cumulative DMFS
Log rank P < 0.02
PLOD2 expression
Low
High
Years
d
ER- HER2-
Cumulative DMFS
Log rank P < 0.01
PLOD2 expression
Low
High
Years
e
ER+ / PR+
HR=1.28 (0.8 - 2.05)
Log rank P = 0.29
HR=1.04 (0.65- 1.65)
Log rank P = 0.88
Probability of Relapse Free Survival
Time (months)
f
HER2+
HR=1.37 (0.75 - 2.5)
Log rank P = 0.3
HR=1.31 (0.72 - 2.39)
Log rank P = 0.37
Time (months)
g
Triple-negative
HR=1.61 (1.05 - 2.48)
Log rank P = 0.029
HR=2.02 (1.3 - 3.12)
Log rank P = 0.00049
PLOD2 Expression
Low
High
Time (months)

**Supplementary Figure 11: PLOD2 correlations with distant metastasis free survival and probability of relapse free survival.** (**a**) Log2 scaled data graphs showing relative PLOD2 gene expression levels in patients stratified by estrogen and epidermal growth factor receptor two (ER/HER2) status (*ER+ HER2- n=314; ER- or + /HER2+ n=73; ER-/HER2- n=133*). Statistical analysis was performed to compare PLOD2 expression levels among subtypes using Kruskal-Wallis test (***p < 0.001) and Mann-Whitney U test for individual comparisons (***p < 0.001). (**b**) Line graphs showing distant metastasis-free survival (DMFS) for patients with estrogen receptor positive and epidermal growth factor receptor two negative breast tumors (ER+/- HER2-; *low n=157 & high=157*). (**c**) Line graph of DMFS for patients with estrogen receptor negative or negative and epidermal growth factor receptor two positive breast tumors (ER- or +/HER2+; *low n=36 & high=37*). (**d**) Line graph of DMFS for patients with estrogen receptor negative epidermal growth factor receptor negative breast tumors (ER- HER2-; *low n=66 & high=67*). Error bars represent minimum and maximum values for each group. Statistical analyses were performed to compare PLOD2 and DMFS for each subtype or Log-rank (Mantel-Cox) test (**P<0.02* and **P<0.01* for ER+/-HER2+ and TN, respectively). (**e-g**) Kaplan-Meier curves indicating the probability of relapse-free survival assessed in ER+/PR+ (**e**), HER2+ (**f**), and triple negative (**g**) breast cancer patients up to 16 years after diagnosis. A correlation between PLOD2 (gene encoding LH2) expression and RFS has been determined using an online tool (http://kmplot.com/analysis/) as described in the methods. **Top and bottom panels** represent two distinct Affymetrix PLOD2 probes (202619 and 202620) from the same database. For Kaplan-Meier curves, statistical analyses were performed using LogRank test.

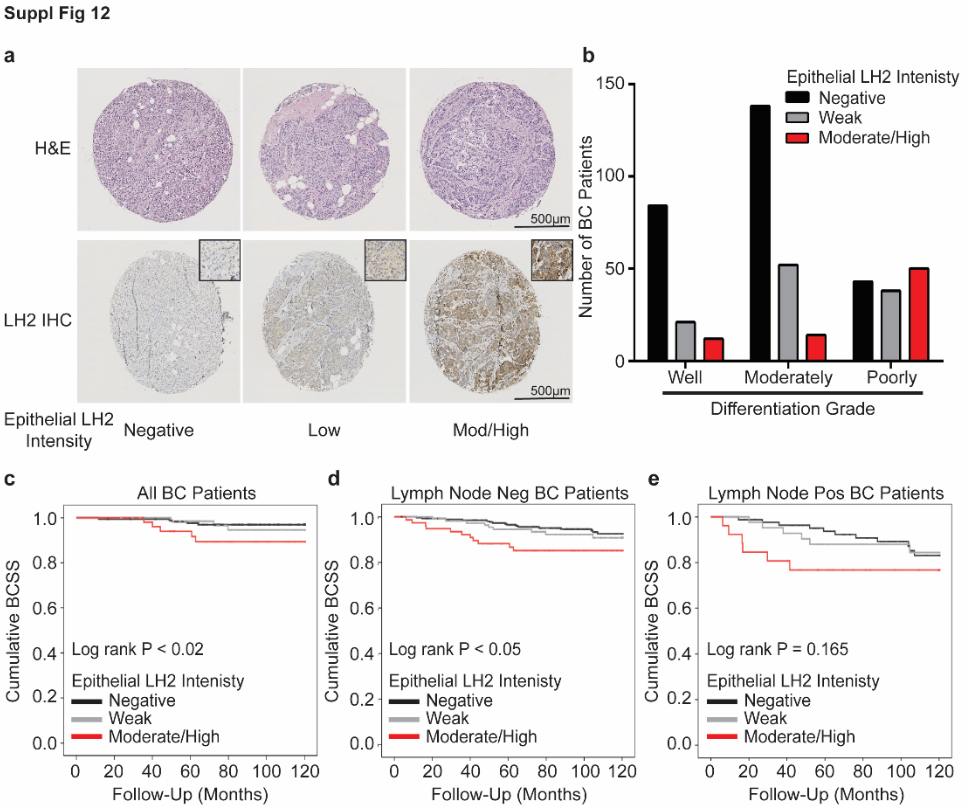

**Supplementary Figure 12: Assessment of neoplastic epithelial LH2 expression in breast cancer.** (**a**) Tumor samples from incident breast cancer cases were collected, and a tissue microarray (TMA) including two 1-mm cores from each tumor was constructed. Neoplastic epithelial LH2 staining was assessed with semi-quantitative intensity score and stratified as negative, low, or moderate/high. (**b**) Clinical correlation of neoplastic epithelial LH2 intensity score with tumor grades. (**c**) Kaplan-Meier curves indicating cumulative breast cancer specific survival (BCSS) based on epithelial LH2 intensity assessed in breast cancer patients up to 10 years after diagnosis (LH2 negative n = 271, weak n = 112, moderate/high n = 77). (**d**) BCSS curves by epithelial LH2 intensity including only axillary lymph node negative patients (LH2 negative n = 175, weak n = 67, moderate/high n = 50). (**e**) BCSS curves by epithelial LH2 intensity including only axillary lymph node positive patients (LH2 negative n = 84, weak n = 42, moderate/high n = 26). For tumor grade and LH2 intensity score, statistical analysis was performed using a linear-by-linear association (***p < 0.001). For Kaplan-Meier curves, statistical analyses were using LogRank test.
